# Epsins oversee smooth muscle cell reprograming by influencing master regulators KLF4 and OCT4

**DOI:** 10.1101/2024.01.08.574714

**Authors:** Beibei Wang, Kui Cui, Bo Zhu, Yunzhou Dong, Donghai Wang, Bandana Singh, Hao Wu, Kathryn Li, Shahram Eisa-Beygi, Yong Sun, Scott Wong, Douglas B. Cowan, Yabing Chen, Mulong Du, Hong Chen

## Abstract

Smooth muscle cells in major arteries play a crucial role in regulating coronary artery disease. Conversion of smooth muscle cells into other adverse cell types in the artery propels the pathogenesis of the disease. Curtailing artery plaque buildup by modulating smooth muscle cell reprograming presents us a new opportunity to thwart coronary artery disease. Here, our report how Epsins, a family of endocytic adaptor proteins oversee the smooth muscle cell reprograming by influencing master regulators OCT4 and KLF4. Using single-cell RNA sequencing, we characterized the phenotype of modulated smooth muscle cells in mouse atherosclerotic plaque and found that smooth muscle cells lacking epsins undergo profound reprogramming into not only beneficial myofibroblasts but also endothelial cells for injury repair of diseased endothelium. Our work lays concrete groundwork to explore an uncharted territory as we show that depleting Epsins bolsters smooth muscle cells reprograming to endothelial cells by augmenting OCT4 activity but restrain them from reprograming to harmful foam cells by destabilizing KLF4, a master regulator of adverse reprograming of smooth muscle cells. Moreover, the expression of Epsins in smooth muscle cells positively correlates with the severity of both human and mouse coronary artery disease. Integrating our scRNA-seq data with human Genome-Wide Association Studies (GWAS) identifies pivotal roles Epsins play in smooth muscle cells in the pathological process leading to coronary artery disease. Our findings reveal a previously unexplored direction for smooth muscle cell phenotypic modulation in the development and progression of coronary artery disease and unveil Epsins and their downstream new targets as promising novel therapeutic targets for mitigating metabolic disorders.

## Introduction

Atherosclerosis is a chronic inflammatory disease characterized by the progressive formation of plaques on the arterial walls that constitutes the primary pathological process for the development and progression of coronary artery disease (CAD)^1,2^. In advanced stage of the disease, rupture of the atherosclerotic plaques causes atherosclerotic thrombosis^3^ leading to life-threatening consequences such as myocardial infarction and stroke^4^. Aortic smooth muscle cells (SMCs) are one of the major cellular components of the atherosclerotic plaques^5^, contributing to the formation of both the fibrous cap and the necrotic core via a process called SMC phenotypic modulation or switching^6^. A paradigm largely built on results from *in vitro* studies seemed to suggest that during atherosclerosis, SMCs migrate into intimal space, proliferate, and transdifferentiate into macrophage-like phenotypes characterized by the expression of *Lgals3* and acquired increased phagocytic activity^7–9^. Those proinflammatory macrophage-adopting SMCs engulf oxidized lipid and dead cells^10^ and eventually become foam cells that enlarge lipid-laden necrotic core and destabilizes plaques^11^. Whereas other studies highlight the conversion of SMCs into synthetic SMC phenotypes which could contribute to the protective fibrous plaque cap and stabilizing plaques^6,8,11^. Utilizing genetic lineage tracing and single cell RNA sequencing techniques, one recent study elucidated that in atherosclerotic lesions SMCs transdifferentiate into a fibroblast-like phenotypes (referred to as myofibroblasts) that express Lgals3 with acquired lipid-phagocytosis capacity. Very few, if any, SMC-derived macrophage like cells were identified in this study^12^. Those myofibroblasts strengthen the fibrous cap and therefore stabilize the plaques^12^. Whereas using a similar approach, another study identified the emergence of a population of stem, endothelial and monocyte/macrophage (SEM) lineage cells derived from SMC in response to atherosclerotic stimuli in addition to significant number of SMC-derived macrophage like cells in the atherosclerotic lesions^13^. Those SMC-derived SEM cells have the potential to differentiate further into macrophages, fibrochondrocytes as well as SMCs^13^. While the SEM lineage of cells express the endothelial marker Vcam-1, the potency of those cells to differentiate into endothelial cells (ECs) to participate in endothelial repair in atherosclerotic lesions has not been investigated.

Krüppel-like factor 4 (KLF4) is a zinc finger transcription factor that plays a critical role in cell fate decision^14^. Numerous studies have shown the arthero-prone function of KLF4 in SMCs^7,15^. KLF4 controls SMC phenotypic plasticity by suppressing the expressions of SMC markers *Acta2*, *Tagln*, *Myh11*, and *Cnn1*^7,16^. Interestingly, the stability of KLF4 is regulated by posttranslational modifications such as methylation and ubiquitination. VHL3-mediated ubiquitination promotes KLF4 degradation by proteosomes whereas methylation inhibits KLF4 ubiquitination, therefore enhances KLF4 protein level^17^. In sharp contrast to KLF4, the stem cell pluripotent transcription factor OCT4 in SMCs is athero-protective in that SMC-specific deletion of OCT4 led to increased size of necrotic core and decreased fibrous plaque cap, and therefore destabilized plaques^18^. A recent study further demonstrated that OCT4 is activated and inhibits intima formation after vascular injury^19^. Those phenotypes in OCT4- SMC knockout mice are exactly the opposite to KLF4-SMC knockout mice^7^. Further studies employing chromatin immunoprecipitation and sequencing (chipseq) identified that KLF4 and OCT4 control nearly opposite patterns of gene expression in SMC^15^. How the counteracting function of KLF4 and OCT4 is coordinated during SMC phenotypic switching remains unclear. The Epsin family of endocytic adaptors including Epsin1 and Epsin2 plays an essential role in the pathology of atherosclerosis via regulating endothelial-to-mesenchymal transition of ECs, as well as lipid uptake, cholesterol efflux, and IP3R1 degradation in macrophages within the atherosclerotic lesions^20–22^. Nevertheless, whether SMC-intrinsic Epsins contribute to the regulation of SMC plasticity, and atherosclerotic pathogenesis has not been explored. Recent studies suggest that Epsins are crucial for transforming growth factor (TGF)-beta receptor endocytosis and signaling^20^ in ECs. TGF-beta activates the *Acta2* (encoding αSMA) expression through promoting KLF4 degradation^23,24^. However, whether Epsins control SMC phenotypic switching through regulating KLF4 stability has not been investigated.

In this study, we profiled cellular components of aortae isolated from atherosclerotic mice at the single-cell level and explored the role of SMC-intrinsic Epsins in the pathogenesis of atherosclerosis. We identified a unique cluster of endothelial-like cells transdifferentiated from SMCs in the atherosclerotic aortae. Those endothelial-like cells can integrate into intima and participate in the repair of endothelial injury caused by atherosclerotic stimuli. *In vitro*, those ECs are equipped with acetylated-low density lipoprotein (ac-LDL) endocytosis capacity. Loss of expression of Epsins in SMCs, on one hand, enhanced KLF4 degradation and therefore promoted SMC marker gene expression. On the other hand, Epsins-deficiency promotes trans- differentiation of SMCs into endothelial-like cells by increasing the protein level of OCT4. At the molecular level, Epsins stabilize KLF4 through inhibiting ubiquitinated KLF4 degradation by directly binding to KLF4 through the Epsins’ ENTH+UIM domain. Mice deficient for both Epsins1&2 specifically in SMCs are more resistant to western diet-induced atherosclerosis. Those findings uncovered a novel function that is independent of Epsins’ endocytic activity in promoting the pathogenesis of atherosclerosis.

## STAR METHODS

### Mice

All animal experiments followed institutional guidelines. Mouse protocols were approved by the Institutional Animal Care and Use Committee (IACUC) of Boston Children’s Hospital, MA, United States.

All mice including *ApoE^-/-^* mice (stock#002052, Jackson Research Laboratory) and *Epsin1^fl/fl^;Epsin2^-/-^* mice used are backcrossed to C57BL/6 (stock#00664, Jackson Research Laboratory) genetic background. SMC-specific deletion of Epsin was established by crossing *Epsin1^fl/fl^;Epsin2^-/-^*mice with SMC-specific SMMHC (*Myh11*)*-iCreER^T2^* transgenic mice (stock# 019079, Jackson Research Laboratory)^25^ as *Epsin1^fl/fl^;Epsin2^-/-^/Myh11-iCreER^T2^* mice. *Epsin1^fl/fl^;Epsin2^-/-^/Myh11-iCreER^T2^* mice were further crossed to *ApoE^-/-^* background to generate the compound mutant mouse strain-*Epsin1^fl/fl^;Epsin2^-/-^/Myh11-iCreER^T2^/ApoE^-/-^*. The details of the SMC-specific deletion of Epsin and *ApoE^-/-^* control mice used in this study were described in Figure S1E. *Myh11-iCreER^T2^-*eYFP^stop/fl^ mice were bred to *ApoE^-/-^*mice to generate *Myh11-iCreER^T2^-*eYFP^stop/fl^*/ApoE^-/-^* mice as controls. These mice were further bred to *Epsin1^fl/fl^;Epsin2^-/-^/Myh11-iCreER^T2^/ApoE^-/-^*mice to establish *Epsin1^fl/fl^;Epsin2^-/-^/Myh11- iCreER^T2^-*eYFP^stop/fl^*/ApoE^-/-^* mice. The details of the SMC-lineage tracing mice and control mice were described in Figure S5B. *Myh11-iCreER^T2^* bacterial artificial chromosome transgene is localized on the Y chromosome, so only male mice were used^26^. 10 μg/g body weight of 4- Hydroxytamoxifen were injected intraperitoneally to induce SMC-specific deletion of *Epsins* and to activate the *eYFP* gene expression at 6 to 8 weeks of age. Then, the mice were fed a western diet (D12079B, New Brunswick, USA) for 9-20 weeks.

### Primary Mouse SMCs Isolation

Mice were anesthetized with isoflurane. Thoracic aortas were harvested from mice and placed in 1× Hank’s balanced salt solution (HBSS) supplemented with penicillin and streptomycin (P/S) at 4°C for 1 hr. Vessels were placed in a sterile culture plate and enzyme solution (Collagenase Type I 5 mg/mL + Collagenase Type IV 5 mg/mL + Liberase Blendzyme 3 0.4 U/mL) was added to cover the vessels. The plates were placed in a 37°C incubator for 2 mins. The aortas were transferred to DMEM medium and cut open longitudinally with scissors. The adventitial layer and EC layer were removed, the muscularis layer was incubated at 37°C for approximately 15 mins. Next, the muscularis layer of aortae were dissected into smaller pieces. The aortic pieces were then carefully removed and placed into 4% gelatin coated 6-well plate, covered with sterile 22 × 22 mm cover slip and supplemented with 1 mL complete DMEM medium containing 20% FBS, 1% insulin-transferrin-selenium, 10 ng/mL epidermal growth factor and 1% P/S. Plates were placed in a 37°C, 5% CO2 culture incubator and the cell medium was refreshed every 3-4 days. As cells expanded and started to cover area (approximately 7 days), the cover slips were flipped, and cells were seeded into a new 6-well plate (cell side up) and covered with 2 mL complete DMEM medium. The residual tissue pieces were removed from the plate and cells were refed in original plate. Cells were allowed to continue to grow until they reach confluence. Cells were weaned into 10% FBS complete DMEM media after 3-5 passages, depending on their viability.

### Small interfering RNA (siRNA) Transfection

siRNA transfection was performed according to the manufacturer’s instructions. Briefly, primary SMCs were transfected by RNAiMAX (CAT#13778, Invitrogen) with either scrambled siRNA duplex or Epsin1 (UGCUCUUCUCGGCUCAAACUAAGGG) or Epsin2 siRNA duplexes (AAAUCCAACAGCGUAGUCUGCUGUG) designed by Ambion® Silencer® Select Predesigned siRNAs (Invitrogen), or ON-TARGETplus Mouse Pou5f1 siRNA (CAT#J-046256-05-002, Invitrogen). At 48 hrs post transfection, cells were processed for western blot assays.

### Immunofluorescent Staining

Human samples: All human samples are from Maine Health Institute for Research Biobank, Maine Medical Center, the details of the samples were described previously^22^. Samples were deparaffinized twice in xylene (15 mins for each time), immersed in graded ethanol (100%, 100%, 95%, 90%, 80%, and 70%, each for 3 mins), washed in running tap water. After blocking endogenous peroxidase activity, the samples were blocked in blocking buffer (PBS with 3% donkey serum, 3% BSA and 0.3% Triton X-100), and incubated with the primary antibodies, anti-αSMA and Epsin1, Epsin2, KLF4 or VHL (1:70 to 1:300 dilution in blocking buffer), 4°C overnight. Respective secondary antibodies conjugated to fluorescent labels (Alexa Flour 488 or 594; 1:200) were added for 2 hrs at room temperature. The sections were mounted with Fluoroshield^TM^ histology mounting medium with DAPI.

Mouse samples: Mouse aortic root and brachiocephalic trunk cryosections were heated to room temperature for 30 mins, fixed in 4% paraformaldehyde for 15 mins and blocked in blocking buffer for 1 hr. Sections were then incubated with the primary antibodies4°C overnight, followed by incubation with the respective secondary antibodies conjugated to fluorescent for 2 hrs at room temperature. The sections were mounted with Fluoroshield^TM^ histology mounting medium containing DAPI.

Cell staining: SMCs were plated on the 18 mm coverslips and washed with PBS for 3 times, fixed in 4% paraformaldehyde for 15 mins and blocked with blocking buffer for 1 hr. Coverslips were incubated with the primary antibodies, 4°C overnight, followed by incubation with the respective secondary antibodies conjugated to fluorescent labels for 1 hr at room temperature. Antibody list, clones and catalogue numbers used for staining are provided in the Table S1. The sections were mounted with Fluoroshield^TM^ histology mounting medium containing DAPI. Immunofluorescent images were captured using a Zeiss LSM880 confocal microscope and analyzed with ZEN-Lite 2012 software and HIH ImageJ software.

### Atherosclerotic Lesion Characterization

The whole aortae were collected and fixed in 4% paraformaldehyde. Next, the aortas were stained with Oil Red O for *en face* analysis. Hearts and brachiocephalic trunk were embedded in O.C.T and sectioned at 8 microns. Lesion area of the aortic root was quantified by hematoxylin and eosin staining. Neutral lipids deposition was determined by Oil Red O staining. Aortic lesion size and lipid content of each animal were obtained by an average of three sections from the same mouse.

### *En face* Oil Red O Staining

Whole aortae were dissected symmetrically, pinned to parafilm to allow the *en face* exposed and fixed in formalin for 12 hrs. The aortae were washed in PBS for 3 times and rinsed in 100% propylene glycol, followed by staining with 0.5% Oil Red O solution for 20 mins at 65°C. The samples were then put in 85% propylene glycol for 2 mins, followed by three washes in DD Water. Slides were next incubated with hematoxylin for 30 sec, rinsed in running tap water. Imaging was performed using a Nikon SMZ1500 stereomicroscope, SPOT Insight 2Mp Firewire digital camera, and SPOT Software 5.1.

### Oil Red O Staining of Cryostat Section

Cryostat sections of mouse aortic root and brachiocephalic trunk were washed in PBS for 2 mins, then fixed in 4% paraformaldehyde for 5 mins. Slices were washed in PBS followed by staining with freshly prepared 0.5% Oil Red O solution in isopropanol for 10 mins at 37°C. Slices were then put in 60% isopropanol for 30 sec, followed by 3 washes in water. Slices were next incubated with hematoxylin for 30 sec, rinsed in running tap water, and mounted with 90% Glycerin.

### Hematoxylin and Eosin Staining

Cryostat sections of mouse aortic root and brachiocephalic trunk were washed in PBS for 2 mins, then fixed in 4% paraformaldehyde for 5 mins. Next, slides were stained with 0.1% hematoxylin for 2 mins followed by washing under running tap water for 2 mins. Slices were then dipped in Eosin working solution for 20 sec, quickly rinsed with tap water, dehydrated using graded ethanol (95% and 100% ethanol), followed by rendering of samples transparent by incubation in 100% xylene for 1 min. Slices were mounted in synthetic resin.

### Van Gieson’s Staining

Van Gieson’s staining was performed based on manufacturer’s instructions. In brief, Cryostat sections of mouse aortic root and brachiocephalic trunk were washed in PBS for 2 mins, then fixed in 4% paraformaldehyde for 5 mins. Slices were placed in Elastic Stain Solution (5% hematoxylin + 10% ferric chloride + Lugol’s lodine Solution) for 20 mins, then rinsed under running tap water. Then, slices were dipped in differentiating solution 20 times and in sodium thiosulfate solution for 1 min, following with rinsing under running tap water. Slices were dehydrated in 95% and 100% alcohol once, respectively, cleared and mounted in synthetic resin.

### RNA Isolation and Quantitative Real-time PCR

Total RNA was extracted using RNeasy^®^ Mini Kit, based on manufacturer’s instruction. cDNA was synthetized by reverse transcription using the iScript cDNA Synthesis Kit (Bio-Rad Laboratories, CA, United States). Quantitative PCR (qPCR) was performed with specific primers using SYBR^®^ Green PCR Master Mix reagent in StepOnePlus Real-Time PCR System. Cdna-specific primers can be found in Table S2.

### Immunoprecipitation and Western Blotting

For immunoprecipitation, SMCs were lysed with RIPA buffer (50 mM Tris, pH 7.4, with 150 mM NaCl, 1% Nonidet P-40, 0.1% SDS, 0.5% sodium deoxycholic acid, 0.1% sodium dodecyl sulfate, 5 mM N-ethylmaleimide and protease inhibitor cocktail). For KLF4 ubiquitination experiments, SMCs were lysed using denaturing buffer (1% SDS in 50 mM Tris, pH 7.4) and boiled at 95°C for 10 mins to denature protein complexes. Lysates were re-natured using nine volumes of ice-cold RIPA buffer, then prepared for immunoprecipitation as follows: Cell lysates were pre-treated with Protein A/G PLUS-Agarose (sc-2003, Santa Cruz Biotechnology) at 4°C for 2 hrs to remove nonspecific protein, followed by centrifugation at 12000 rpm for 5 mins at 4°C. Supernatant was transferred to a new tube, incubated with Protein A/G PLUS-Agarose and antibodies against Epsin1 or KLF4 or ubiquitin at 4°C overnight. Mouse IgG was used as negative control. Protein A/G beads were washed with RIPA buffer for 2 times, followed by PBS for 1 time. Then, beads were suspended with 80 μL 2× loading buffer and heated at 95°C for 10 mins. After centrifugation, precipitated proteins were visualized by Western blot. Proteins were resolved by SDS-PAGE gel and electroblotted to nitrocellulose membranes. NC membranes were blocked with 5% milk (w/v) and blotted with antibodies. Western blots were quantified using NIH Image J software.

### Differentiation of SMCs to EC Phenotype

EC function was tested with DiI-ac-LDL Staining Kit based on manufacturer’s instructions. Briefly, SMCs were planted onto 12-mm slides until they reached 95% confluence. Next, the cells were treated with 100 μg/mL oxLDL for 4 days. Then, 10 μg/mL DiI-ac-LDL (CAT#022K, Cell Applications) was added, instead of oxLDL, in the medium onto each 12-mm slide. The slides were placed in a 37°C, 5% CO2 incubator for 6 hrs. The cells were washed 3 times with a wash buffer. The slides were mounted with a cover slip using mounting solution. Images were taken using a Zeiss LSM880 confocal microscope and analyzed with HIH ImageJ software.

### Flow Cytometry and Cell Sorting

Prepare a single cell suspension isolated from eYFP-SMC mice aorta, thoracic aortae were isolated as previously reported. The aortae were cut into small pieces and moved into a new dish containing enzyme solution, incubated at 37°C in the incubator for about 1 hr. After an hour, the plates were washed with 2 mL warmed DMEM medium (DMEM + 5% FBS + P/S). Cells were collected by spinning 1500 rpm for 5 mins. The supernatant was carefully removed and 0.5 mL sterile PBS containing 1% BSA was added. Total eYFP-tagged SMCs were sorted as live by using BD FACSARIA II.

Single-cell suspensions used for intracellular staining were fixed in ice-cold 4% PFA, following treatment with 150 μL permeabilization buffer (1% Triton X-100 in PBS). Next, 1 μg blocking IgG was added and the samples were incubated at room temperature for 15 mins. Intracellular cytokines were stained with antibodies against CD31, VE-Cadherin, KLF4 or OCT4. Total eYFP-tagged SMCs were sorted as live, CD31^+^ or VE-Cadherin^+^ for KLF4 or OCT4 expression in *Epn1&2-SMC^iDKO^/eYFP^+/-^/ApoE^-/-^*and *eYFP^+/-^/ApoE^-/-^* mice. Antibody list, clones and catalogue numbers used for staining were provided in Table S1. BD FACSARIA II was used to collect raw data from flow cytometry experiments. All data files were analyzed using FlowJo version 9.

### Cell Culture and Plasmids Transfection

The HEK 293T cell line (ATCC no. CRL-11268) was used for plasmid transfection to map the binding sites of Epsin to KLF4. Flag-tagged Epsin1WT, Epsin1ΔUIM, Epsin1ΔENTH truncation constructs, and pcDNA vector were prepared previously in our lab. pCX4-KLF4 (Plasmid #36118) were purchased from AddGene. HEK 293T cells were cultured in DMEM (10% FBS and 1% Pen-Strep) at 37°C in humidified air containing 5% CO2 atmosphere and transfected using Lipofectamine 2000 as instructed by the manufacturer.

The primary SMCs isolated from *Epn1&2-SMC^iDKO^/eYFP^+/-^/ApoE^-/-^* were infected with adenovirus (Ad)-KLF4 or Ad-null for 48 hrs^27^. Ad-KLF4 and Ad-null are gifts from Dr. John Y.-J. Shyy, University of California, San Diego.

### Single-cell Preparation and Data Processing

Single cell of mouse arterial specimens was prepared as above. The cell viability exceeded 90% and was determined under the microscope with trypan blue staining. 20 μL of cell suspension was calculated to contain ∼20,000 cells for each sample. Single-cell capturing and library construction were performed using the Chromium Next GEM Single Cell 3’ Reagent Kits v3.1 (10× Genomics) according to the manufacturer’s instructions. In brief, 50 μL of barcoded gel beads, 45 μL partitioning oil and 70 μL cell suspension were loaded onto the Chromium Next GEM Chip G to generate single-cell gel beads-in-emulsion. Captured cells were lysed and the transcripts were reverse-transcribed inside individual gel beads-in-emulsion. Full-length cDNA along with cell barcodes were amplified via PCR. The sequencing libraries were constructed by using the 3’ Library Kits. Each sample was processed independently. The constructed libraries were sequenced on an Illumina NovaSeq platform.

Similar to the method employed in our previous study^20^, raw sequencing data of FASTQ files were processed using CellRanger (version 3.0.2, 10× Genomics) with default parameters and mapped to mouse reference genome mm10, as well as annotated via a Ensembl 93 annotation, to generate matrices of gene counts by cell barcodes. We used *Seurat* package^28^ to conduct quality controls and downstream analyses. For quality controls, genes expressed in less than 10 cells and cells with less than 100 genes were initially removed from the datasets. The subsequent filters at the cell level met the following criteria of number of genes detected per cell > 250, number of UMIs per cell > 500, log10 transformed number of genes detected per UMI > 0.8, and mitochondrial counts ratio < 0.2. Raw unique molecular identifier (UMI) counts were normalized and regressed by mitochondrial mapping percentage using *SCTransform* function. Possible batch effects derived from different conditions on mouse models were adjusted using *Harmony* package^29^. Dimension reduction was performed using principal-component analysis (PCA) with *RunPCA* function. Two-dimensional Uniform Manifold Approximation and Projection (UMAP) was used for visualization. Graph-based clustering was performed on the integrated dataset with a default method of K-nearest neighbor (KNN) graph. Cell clusters were identified using the graph observed above with a resolution parameter ranging from 0.1 to 1.2. In this study, we divided cells into 26 clusters underlying the resolution parameter of 0.8, which were further grouped into 10 cell subgroups.

### Cleavage Under Targets and Tagmentation (CUT&Tag)

SMCs isolated from aortae of both *ApoE^-/-^* and *Epn1&2-SMC^iDKO^/ApoE^-/-^*mice (n=3) were subjected for CUT&Tag assay protocol according Henikoff’s lab with minor modification^30^. Briefly, 100,000 isolated VSMCs were bund to Concanavalin-A-coated beads, and then bind primary antibodies of KLF4 and OCT4 (1:100 dilution) to the cell and Concanavalin A-coated beads complex for overnight at 4 °C. The pig-anti-rabbit secondary antibody (1:100 dilution) was added into the mixture above for 1 hr incubation. After secondary antibody incubation, the mixture was washed by Dig-wash buffer twice, bind pAG-Tn5 adapter complex for 1 hr, and then washed by Dig-300 buffer twice. The complex mixture was incubated at 37°C for tagmentation for 1 hr. After tagmentation, DNA fragments were extracted and further for PCR and post-PCR clean-up. Finally, the established libraries were sent for DNA sequencing.

We followed the pipeline https://yezhengstat.github.io/CUTTag_tutorial to analyze the CUT&Tag data. Briefly, sequence reads that passed the quality control by FastQC were aligned to the mm10 mouse reference genome using Bowtie2. Peak calling was performed by *SEACR* R package (PMID: 31300027), which provided enriched regions from chromatin profiling data. *DESeq2* R package (PMID: 25516281) was used to analyze the differential enriched peaks of each transcription factor between *ApoE^-/-^* and *Epn1&2-SMC^iDKO^/ApoE^-/-^* mice. The target genes harboring the differential peaks were used for further signature score calculation in scRNA data.

### Trajectory Analysis

To calculate the RNA velocity of the single cells, we used the CellRanger output BAM file and GENCODE file to together with the velocyto^31^ CLI v.0.17.17 to generate a loom file containing the quantification of spliced and unspliced RNA. Next, we built a manifold, clustered the cells and visualized the RNA velocities using *scVelo*^32^. cytoTRACE analysis with default parameter^33^ was performed to predict differentiation states from scRNA-seq data based on the simple yet robust observation that transcriptional diversity decreases during differentiation, to complement the trajectory analysis from RNA velocity. Pseudotime was analyzed using *Monocle* package^34^ with reduceDimension and plot_cell_trajectory functions.

### Cellular Interactions among Different Cell Types

To describe potential cell-to-cell communications, we leveraged the *CellChat* R package^35^ to infer the cellular interactions based on the normalized scRNA-seq dataset. The algorithm of CellChat could examine the ligandreceptor interactions significance among different types of cells based on the expression of soluble agonist, soluble antagonist, and stimulatory and inhibitory membrane-bound co-receptors. By summing the probabilities of the ligand-receptor interactions among a given signaling pathway, we could calculate the communication probability for the pathway. In brief, we followed the official workflow and loaded the normalized counts into CellChat and applied the preprocessing functions *identifyOverExpressedGenes*, *identifyOverExpressedInteractions* and *projectData* with standard parameters set. As database we selected the ‘Secreted Signaling’ pathways and used the pre-compiled ‘Protein-Protein-Interactions’ as a priori network information. For the main analyses the core functions *computeCommunProb*, *computeCommunProbPathway* and *aggregateNet* were applied using standard parameters and fixed randomization seeds. Finally, to determine the senders and receivers in the network the function *netAnalysis_signalingRole* was applied on the ‘netP’ data slot.

### Gene-based Genetic Association Analysis

We used the publicly available GWAS summary statistics of CAD in European populations^36^ from a meta-analysis of three datasets, including UK Biobank SOFT CAD GWAS, the CARDIoGRAMplusC4D 1000 Genomes-based GWAS^37^, and the Myocardial Infarction Genetics and CARDIoGRAM Exome^38^. The SOFT CAD phenotype in UK Biobank^36^ encompasses individuals with fatal or nonfatal myocardial infarction (MI), percutaneous transluminal coronary angioplasty (PTCA) or coronary artery bypass grafting (CABG), chronic ischemic heart disease (IHD) and angina. CARDIoGRAMplusC4D 1000 Genomes-based GWAS^37^ is a meta-analysis of GWAS studies of mainly European, South Asian, and East Asian, involving 60,801 CAD cases and 123,504 controls. Myocardial Infarction Genetics and CARDIoGRAM Exome^38^ is a meta-analysis of Exome-chip studies of European descent involving 42,335 patients and 78,240 controls.

A total of 8,908,875 SNPs without exome chip data were retained. We extracted SNPs available in individuals of Utah residents (CEPH) with Northern and Western European ancestry from 1000 Genomes Project (Phase I, version 3), and then performed quality control using the following criteria: minor allele frequency (MAF) > 0.01, call rate ≥ 95% and *P* value of Hardy-Weinberg equilibrium (HWE) > 0.01. Eventually, a total of 7,580,209 SNPs were included for further gene analysis. After variant annotation, SNPs were mapped into 17,910 protein-coding genes including the body of the gene or its extended regions (± 20 kb downstream or upstream). The SNP-based *P* values from the GWAS meta-analysis were used as input for the gene-based analysis computed by leveraging a multivariant converging regression model in the Multi-marker Analysis of GenoMic Annotation (MAGMA)^39^. Stringent Bonferroni correction was applied for multiple testing with the genome-wide significance at *P* = 2.79E-6 (0.05/17,910), which generated 68 candidate CAD susceptibility genes for further signature score analysis and pathway enrichment analysis of Gene Ontology by *clusterProfiler* R package.

### Gene signature score calculation

We calculated signature scores on the basis of scRNA data underlying *PercentageFeatureSet* function in Seurat. CUT&Tag signature score of OCT4 and KLF4 were calculated based on the genes harboring the differential peaks CAD GWAS signature score was calculated based on the expression of 68 CAD susceptibility genes, as well as increased and decreased signature score upon the literature review for 68 genes.

### Mendelian Randomization

Summary-level statistics of aptamer-based plasma protein KLF4 were extracted from a large-scale protein quantitative trait loci (pQTL) study in 35,559 Icelanders at deCODE. The levels of protein were rank-inverse normal transformed and adjusted for age and sex. Details on the GWAS can be found in the original publication^40^. Mendelian Randomization (MR) is an analytical method, which uses genetic variants as instrumental variables (IVs) to assess the causal effect of specific phenotypes on outcomes^41^. We performed two-sample MR analysis to obtain causal estimates of plasma protein KLF4 on CAD using the *TwoSampleMR* package^42^. Independent SNPs (LD r^2^ < 0.001, within 10,000 kb) at *P* < 5e-8 were retained as instrumental variables. Inverse-variance-weighted (IVW), weighted median, and MR-Egger regression were primarily used to calculate effect size (β) and corresponding standard error (SE). Heterogeneity was estimated by MR-Egger and IVW methods to assess whether a genetic variant’s effect on outcome was proportional to its effect on exposure. Directional pleiotropy was estimated via MR-Egger intercept test for the presence of horizontal pleiotropy.

### Statistical Analysis

All wet bench data were expressed as mean ± SEM and the statistical analyses were performed with SPSS 16.0. The 2-tailed Student’s *t* test was used for parametric data analyses, ANOVA was used to compare the difference between multiple groups. *P* < 0.05 was considered to be statistically significant.

## Results

### Upregulated Expression of Epsins in VSMCs in Response to Atherosclerotic Stimuli

To explore whether Epsins in SMCs contribute to the pathogenesis of atherosclerosis, we examined Epsins expressions in atherosclerotic lesions from patients with various disease burdens. In human coronary arteries with disease histologically classified as no lesions, mild lesion with small plaques, and severe lesions with large plaques, we observed that Epsin1 and Epsin2 were expressed in SMCs and in the atherosclerotic lesions. Importantly, the expression of Epsins 1&2 seemed to be enhanced with the increase of the severity of the disease. (Figure S1A).

To evaluate Epsins expression in SMCs in mouse atherosclerotic plaques, we compared Epsins expression in *ApoE^-/-^* mice fed on normal or western diet for 16 weeks. We found that Epsin1 expression was dramatically increased in the plaques of mice fed on western diet compared to those from mice fed on normal diet (Figure S1B). We next assessed *Epsins* transcript abundance in primary SMCs isolated from *ApoE^-/-^* mice and found that treatment of SMCs with oxLDL resulted in a 1.9- and 1.7-fold increase in *Epsin1* and *Epsin2* transcripts, respectively (Figure S1C,D). Together, these observations indicated atherosclerotic stimuli increases Epsins expression in SMCs both *in vitro* and in atherosclerotic plaques *in vivo*.

### Single-cell Transcriptomics Identified a Novel Cluster of SMC-modulated ECs in the Atherosclerotic Aortae

To determine the role of Epsins in atherosclerosis, we crossed *Epn1^fl/fl^*, *Epn2^-/-^* mice with *Myh11-iCre^ERT2^* transgenic mice with a tamoxifen-inducible iCre recombinase knocked into the SMC-specific *Myh11* locus on a bacterial artificial chromosome^26^. We named the resultant *Epn1^fl/fl^*, *Epn2^-/-^*, *Myh11-iCre^ERT2^* strain as *Epn1&2-SMC^iDKO^* mice. *Epn1&2-SMC^iDKO^* mice were further crossed to *ApoE^-/-^*background (*Epn1&2-SMC^iDKO^/ApoE^-/-^*), injected with tamoxifen to induce the deletion of Epsin1 in SMC (Figure S1E) at the age of 8 weeks, followed by feeding on western diet for 6, 12, and 16 weeks. Immunostaining of aorta sections demonstrated the abrogation of Epsins1&2 in SMCs of aortae from *Epn1&2-SMC^iDKO^*mice after tamoxifen injection (Figure S1F).

We next performed single-cell RNA sequencing on cells isolated from the whole aortae from *ApoE^-/-^ and Epn1&2-SMC^iDKO^/ApoE^-/-^*mice at baseline and those fed western diet for 6, 12 and 16 weeks (Figure 1A). After stringent quality control of scRNA-seq data processing, a total of 151,944 cells across week feeding groups (Figure S2A,B), i.e., *ApoE^-/-^*: normal diet (n = 23,709), 6-week western diet (n = 6,318), 12-week western diet (n = 27,569), and 16-week western diet (n = 22,258); *Epn1&2-SMC^iDKO^/ApoE^-/-^*: normal diet (n = 16,036), 6-week western diet (n = 21,093), 12-week western diet (n = 15,295), and 16-week western diet (n = 19,666) were retained for downstream analysis. Graph-based clustering of the individual datasets visualized by UMAP^43^ and canonical cell marker annotation gave rise to eight main cell clusters (Figure1B, Figure S2C,D), including SMC, modulated SMC (modSMC), modulated SMC with EC markers (modSMC_EC), modulated SMC with fibroblast markers (modSMC_myofibroblast), fibroblast, EC, macrophages, and immune cells.

**Figure 1.**
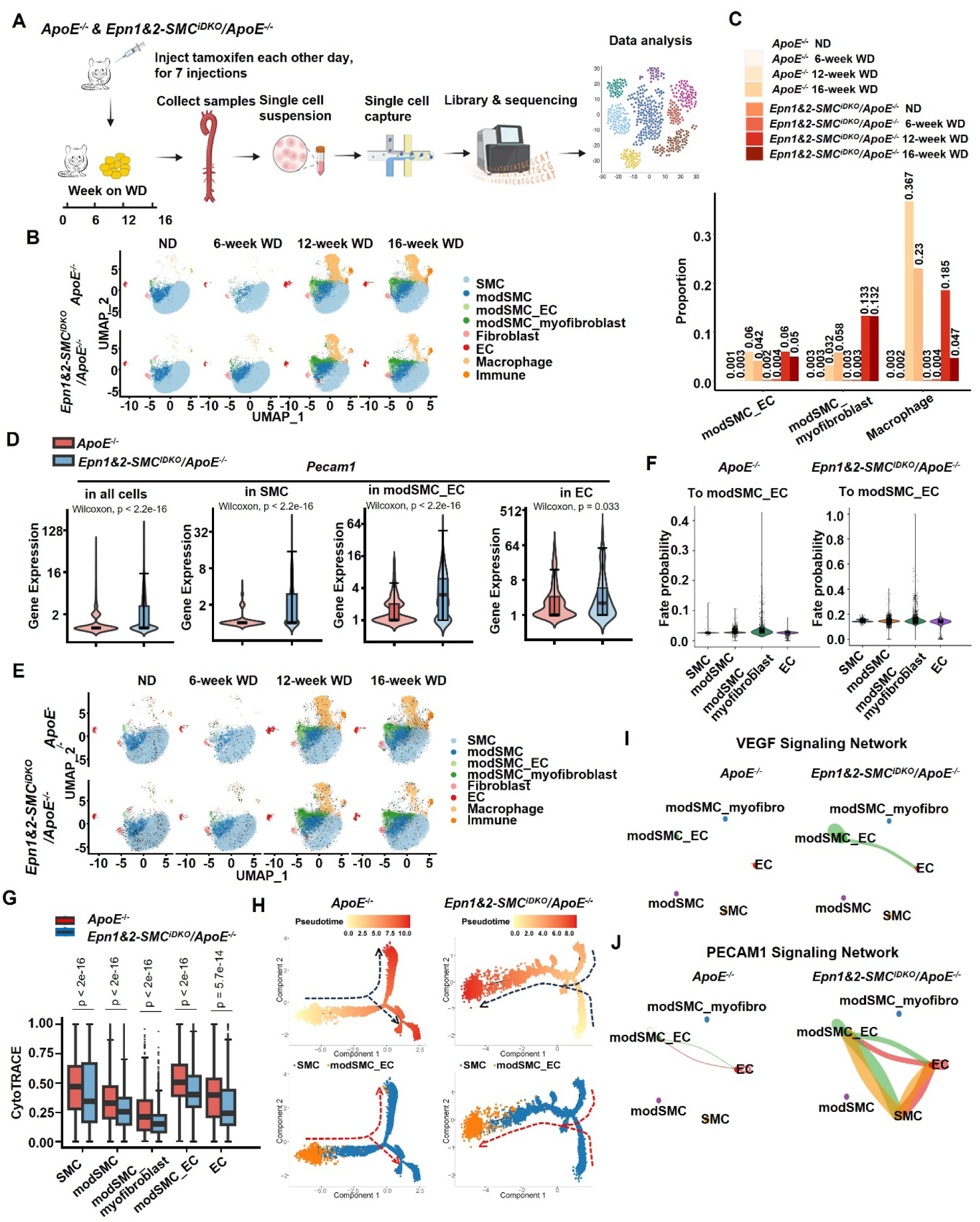
Single-cell Transcriptomic Profiling of Aortae from *ApoE*^-/-^ and *Epn1&2- SMC^iDKO^/ApoE^-/-^* Mice and Cell Transdifferentiation. (A) Mouse model construction and corresponding scRNA-seq experimental workflow. (B) Uniform Manifold Approximation and Projection (UMAP) visualization of eight major cell types of mouse aortae across varying lengths of feeding on ND or WD. Dots represent individual cells, and colors represent different cell populations. (C) Proportion of major cell types. (D) Differential gene expression *Pecam1* in various cell clusters from aortae of *ApoE^-/-^* and *Epn1&2-SMC^iDKO^/ApoE^-/-^* mice. *P* value was calculated by Wilcoxon rank sum test. (E) UMAP visualization of inferred RNA velocity for eight major aortae cell clusters. (F) Fate probability of major cell types transitioning into the modSMC_EC cluster in the aortae of *ApoE*^-/-^ and *Epn1&2-SMC^iDKO^/ApoE^-/-^* mice inferred by *CellRank*. g, CytoTRACE scores of major cell types in aortae of *ApoE^-/-^* and *Epn1&2-SMC^iDKO^/ApoE^-/-^*mice with long-term WD feeding. (H) The trajectory path from SMC to modSMC_EC cells in aortae of *ApoE^-/-^*and *Epn1&2-SMC^iDKO^/ApoE^-/-^* mice inferred by *Monocle2*. The trajectory direction was determined by the predicted pseudotime. The trajectories were colored by pseudotime (up) and cluster identities (down). (I-J) Cell communication networks of VEGF (I) and PECAM1 (J) signaling among major cell clusters in the aortae of *ApoE^-/-^* and *Epn1&2-SMC^iDKO^/ApoE^-/-^*mice calculated using *CellChat*. WD, western diet; ND, normal diet. SMC, aortic smooth muscle cell; modSMC, modulated SMC; EC, endothelial cell.

Of particular interest in our scRNAseq result is the emergence of a cell population that retained conventional SMC markers while also expressed the canonical endothelial marker Pecam1 (Figure S2C). As described above, we defined such a cell population as modSMC_EC. It is of note that the abundance of modSMC_EC increased from negligible in normal diet-fed mice aortae to a significant portion in both and *ApoE^-/-^*and *Epn1&2-SMC^iDKO^/ApoE^-/-^* mice (Figure 1C, Figure S2E). More importantly, SMC-specific deficiency of Epsins led to increased expression of the conventional EC marker Pecam1 in all cell clusters including modSMC_ECs from aortae of *Epn1&2-SMC^iDKO^/ApoE^-/-^* mice compared to that from *ApoE^-/-^* mice (Figure 1D). This observation suggested that atherosclerotic stimuli increased the transition of SMC into modSMC_EC population and Epsins are negative regulators of such a transition. However, the role of the modSMC_ECs which are originated from VSMCs, in the pathogenesis of atherosclerosis is not clear.

### SMC-intrinsic Epsins Inhibits Transdifferentiation of SMCs into ModSMC_ECs in Atherosclerotic Aortae

To investigate the cellular dynamics during the pathological development of atherosclerosis, we performed unsupervised trajectory analysis on the scRNAseq results using RNA velocity algorithms^31^. Cell population trajectory showed little difference between the *ApoE^-/-^* and *Epn1&2-SMC^iDKO^/ApoE^-/-^*mice aortae when fed on normal diet. However, such trajectory changed in aortae from mice feed on western diet for 12 or 16 weeks. The velocity flow of VSMCs toward atheroprone macrophages is decreased with concomitant increased VSMC velocity toward modulated SMCs including modSMC_myofibroblast and modSMC_ECs in the aortae of *Epn1&2-SMC^iDKO^/ApoE^-/-^*mice compared to that in *ApoE^-/-^* mice (Figure 1E). CellRank^44^ analysis of the cell dynamics revealed that multiple cell clusters had increased probability to transit into modSMC_EC population in aortae from *Epn1&2- SMC^iDKO^/ApoE^-/-^*mice compared to that in *ApoE^-/-^* mice (Figure 1F). Taken together with the findings that the proportion of macrophages were drastically decreased in aortae from *Epn1&2- SMC^iDKO^/ApoE^-/-^* mice compared to that in *ApoE^-/-^*mice fed on western diet (Figure 1B,C), those observations suggest that VSMC-intrinsic Epsins promote the recruitment and accumulation of inflammatory macrophages in atherosclerotic lesions while inhibiting the phenotype transition of VSMCs into athroprotective modSMC_myofibroblast and the modSMC_EC population. As the transition was much more notable in the aortae from mice fed on western diet for 12 and 16 weeks (Figure 1E), with disease progression (Figure S1A,B), we will focus on these groups of mice for the rest of our data analysis.

Unsupervised cytoTRACE analysis^33^ is used to predicate the cellular differentiation status.

Cells with low cytoTRACE score indicated a more differentiated status and vice versa. Similar to the velocity analysis, we observed that the predicted cytoTRACE score of transition from SMC to modSMC relevant cells were significantly lower in *Epn1&2-SMC^iDKO^/ApoE^-/-^*mice compared with that in *ApoE^-/-^* mice, indicating a higher probability of transition of SMC lineages into modSMC_EC and modSMC_myofibroblsts in the absence of Epsins (Figure 1G). To further investigate the role of Epsins in the transition of SMC into modSMC_ECs, we performed an unsupervised pseudotime analysis^45,46^ focusing on the transition from SMC to modSMC_EC. In mice fed on western diet for 16 weeks, modSMC_EC served as an intermediate that tends to be converted back into SMC in the aortae of *ApoE^-/-^* mice as it was located at a tree node at the start of pseudotime. Whereas modSMC_EC in the aortae of *Epn1&2-SMC^iDKO^/ApoE^-/-^* was at the end of pseudotime derived from SMC, indicating a more differentiated stage toward EC (Figure 1H). Taken together, those scRNAseq data analysis support the conclusion that VSMC-intrinsic Epsins inhibits SMC transition into modSMC_EC under atherosclerotic conditions.

Cell-to-cell communications analysis using CellChat^35^ revealed increased number and strength of inferred cell-cell interactions in aortae of *Epn1&2-SMC^iDKO^/ApoE^-/-^*compared to that of *ApoE^-/-^* mice (Figure S3A,B). Among the strengthened communications are the ones involved in the transition of other lineage cells into EC such as VEGF-VEGFR^47^, PECAM1 (Figure 1I,J), NOTCH1^48^ and EGF^49^ (Figure S3B-D). To verify those findings in scRNAseq analysis, we performed reverse transcript quantitative PCR (qRT-PCR) and Western blot analysis on homogenates of aortae from tamoxifen injected *ApoE^-/-^* and *Epn1&2- SMC^iDKO^/ApoE^-/-^*mice fed on western diet for 16 weeks. VSMC-specific deficiency of Epsins led to an increase of the EC marker CD31 in aortae from *Epn1&2-SMC^iDKO^/ApoE^-/-^*mice compared to that from *ApoE^-/-^* mice (Figure S3E, Figure 3A). We concluded that Epsins in the VSMCs inhibits signaling flow between cells that promotes the transition of SMCs into endothelial-like cells.

### SMC-intrinsic Epsins Promoted Atherosclerosis through Destabilizing SMCs’ Contractile Phenotype

We next performed Gene Ontology pathway enrichment of differentially expressed genes in SMCs from aortae from *ApoE^-/-^* and Epn1&2-SMCiDKO/*ApoE*^-/-^ mice across normal diet and western diet-fed (Table S5), and notably observed that the presumably atheroprone signaling pathways such as TLR50, ERK^51^, PI3K^52^, TGFβ^20^, NF-κB^53^ were decreased in aortae from *Epn1&2-SMC^iDKO^/ApoE^-/-^* mice compared to that from ApoE-/- mice (Figure 2A). In addition to the compromised atheroprone pathways, cholesterol storage and foam cell differentiation were also decreased in Epsins-deficient VSMCs, suggesting that Epsins in VSMC promoted cholesterol accumulation and foam cell formation which promotes the formation and growth of atherosclerotic plaques. Moreover, Epsins in VSMC promoted the activation of proinflammatory cells and cytokinesis production. All these data underpin an atheroprone role of Epsins in VSMCs that underlies the pathogenesis of atherosclerosis.

**Figure 2.**
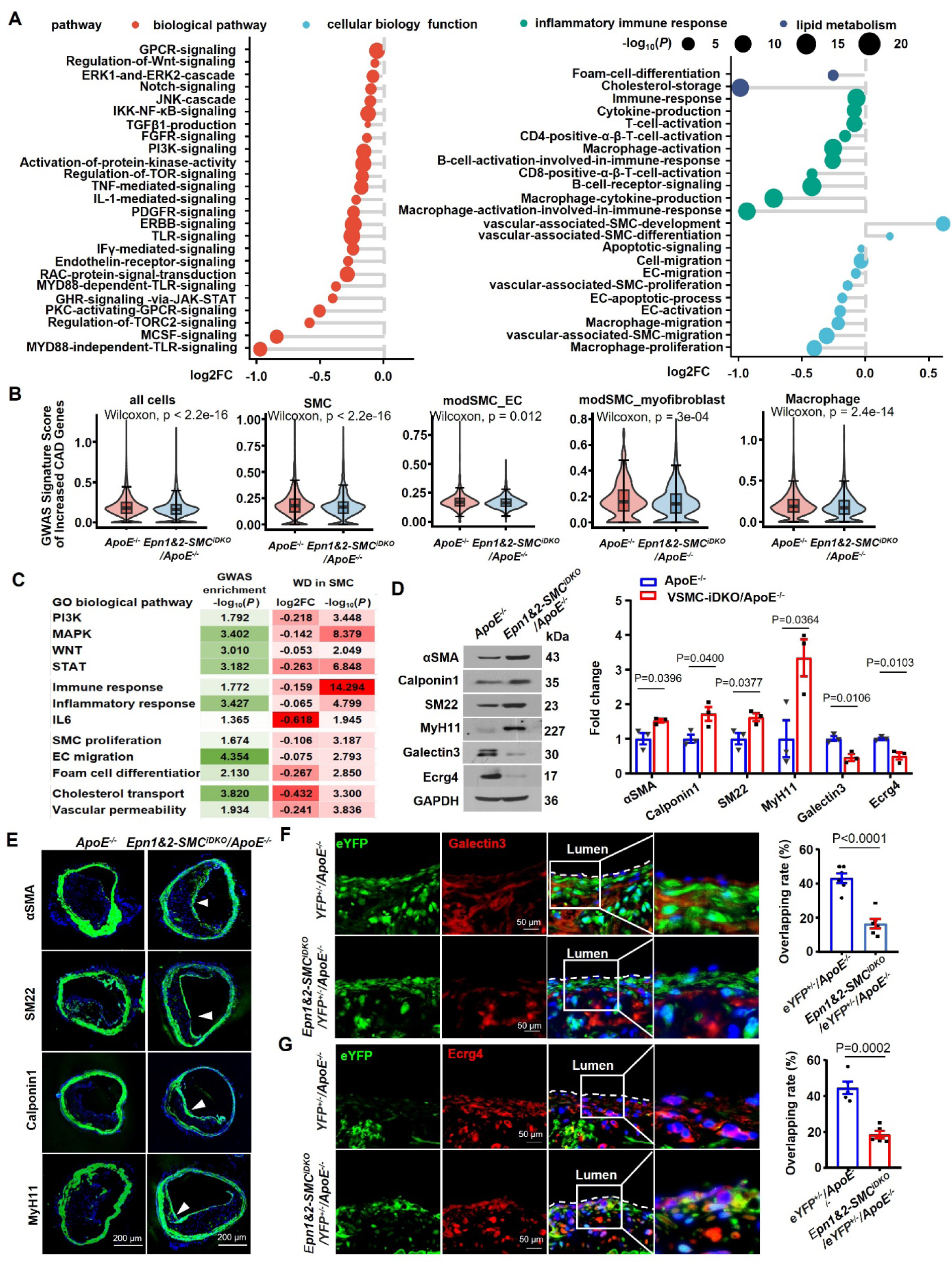
SMC Epsins Destabilize SMC Contraction Phenotype and Promote Atherosclerosis. (A) GO functional annotation and pathway enrichment analysis of differentially expressed genes of scRNAseq data of aortae cells from *ApoE^-/-^* and *Epn1&2-SMC^iDKO^/ApoE^-/-^*mice. Each pathway was scored using *UCell* method deposited in *irGSEA* and *P* value was calculated using Student’s *t*-test. FC, fold change of pathway score by comparing *Epn1&2-SMC^iDKO^/ApoE^-/-^*mice to *ApoE^-/-^* mice. (B) Combined gene expression score of the 19 CAD signature genes in the scRNAseq dataset of aortae from *ApoE^-/-^* and *Epn1&2-SMC^iDKO^/ApoE^-/-^* mice across major cell types. Those 19 genes are associated with increased risk of CAD identified by GWAS of human patients as described in the text. *P* value was calculated by Wilcoxon rank sum test. (C) Enrichment scores of representative atheroprone pathways reveald by GO pathway enrichment analysis on both the 68 CAD signature genes from human GWAS and genes differentially expressed in scRNAseq data of aortae from *ApoE^-/-^* and *Epn1&2-SMC^iDKO^/ApoE^-/-^*mice across normal and long-term WD feeding groups. *P* value was calculated using Student’s *t*-test. FC, fold change of pathway score by comparing *Epn1&2-SMC^iDKO^/ApoE^-/-^* mice to *ApoE^-/-^* mice. (D) Immunoblot of VSMC markers (αSMA, Calponin1, SM22, MyH11), macrophage marker (Galectin3) and fibroblast marker (Ecrg4) in the homogenates of aortae from *ApoE^-/-^* and *Epn1&2-SMC^iDKO^/ApoE^-/-^* mice fed a WD for 16 weeks. All *P* values were calculated using two-tailed unpaired Student’s *t*-test. Data are mean ± s.d. n = 3 independent repeats. e, Immunofluorescence staining for α-SMA, SM22, Calponin1, and MyH11 in brachiocephalic trunk of *ApoE^-/-^*and *Epn1&2-SMC^iDKO^/ApoE^-/-^* mice fed on WD for 16 weeks. Scale bar=200 μm. White arrowhead indicate SMC markers staining on fibrous cap. (F-G) Immunofluorescence staining of macrophage marker Galectin3 (F) and fibroblast marker Ecrg4 (G) in brachiocephalic trunk of YFP-tagged SMC-lineage tracing mice. Scale bar=50 μm. n=5-6 mice. SMC, aortic smooth muscle cell; EC, endothelial cell; GO, Gene Ontology; GWAS, genome-wide associated studie; CAD, coronary artery disease; WD, western diet. (D-G) All *P* values were calculated using two-tailed unpaired Student’s *t*-test. Data are mean ± s.d.

To explore the relevance of our mice scRNAseq data to human diseases, we retrieved 68 CAD signature genes identified through GWAS analysis in European populations^36^ (Figure S4A, Table S3) and mapped the transcript abundance of those genes in our scRNAseq dataset. Overall, the expression of the 68 CAD signature genes are significantly lower in cells across all the population in aortae of *Epn1&2-SMC^iDKO^/ApoE^-/-^* that that from *ApoE^-/-^*mice (Figure S4B). Notably, 19 of 68 CAD genes are associated with increased CAD risk, while 30 are associated with decreased CAD risk (Table S3) in the GWAS dataset. Intriguingly, we observed that the 19-gene cohort associated with increased CAD risk are downregulated in multiple cell types from the aortae of *Epn1&2-SMC^iDKO^/ApoE^-/-^*mice compared to that from *ApoE^-/-^* mice (Figure 2B), while no significant differential expression were found in the 30-gene cohort associated with decreased CAD risk among the two genotypes (Figure S4C).

We next performed pathway enrichment analysis on the 68 CAD signature genes (Table S4). We observed the enrichment of several atheroprone biological process pathways such as PI3K, MAPK, WNT, and STAT, as well as pathways involving inflammatory immune responses and affecting cell phenotypes and functions (e.g., SMC proliferation, EC migration, and vascular permeability) (Figure 2C). Intriguingly, those presumably atheroprone pathways such as PI3K^52^, MAPK^54^, WNT^55^, and STAT^56^, inflammatory immune responses, and cellular function and biology were also identified in pathway enrichment assay using the differentially expressed genes of our scRNAseq results of mouse aortae (Figure 2C). More importantly, those presumably atheroprone pathways are downregulated in aortae SMCs from *Epn1&2- SMC^iDKO^/ApoE^-/-^*mice compared to that from *ApoE^-/-^* mice. Together, combining the GWAS of human data and our scRNAseq data, we conclude that VSMC-intrinsic Epsins are atheroprone.

To explore the mechanism by which Epsins contribute to the pathology of atherosclerosis as suggested by combined data of scRNAseq and GWAS, we harvested aortae from tamoxifen injected *ApoE^-/-^*and *Epn1&2-SMC^iDKO^/ApoE^-/-^* mice fed on western diet for 16 weeks. qRT-PCR analysis showed that the expression of VSMC markers related to VSMC contractile phenotype (i.e., *Acta2*, *Cnn1*, *Tagln*, and *Myh11*)^8^ were significantly higher in the aortae from *Epn1&2-SMC^iDKO^/ApoE^-/-^*mice than that from *ApoE^-/-^* mice (Figure S5A). Western blot on aortae homogenates and immunostaining on aortae sections further showed that VSMC-specific deficiency of Epsins led to stabilization of SMC contractile markers, as well as a decrease of macrophage marker Galectin3 in aortae from *Epn1&2-SMC^iDKO^/ApoE^-/-^*mice compared to that from *ApoE^-/-^* mice (Figure 2D,E). To track VSMC phenotypic switching in atherogenesis, we crossed *Epn1&2-SMC^iDKO^/ApoE^-/-^* mice with Rosa26Stop-floxed eYFP reporter stain of mice^57^ generating in *Epn1&2-SMC^iDKO^/eYFP^+/-^/ApoE^-/-^*compound mutant mice (Figure S5B). When those mice were injected with tamoxifen at the age of 8 weeks, all SMCs and cells derived from SMCs subsequently will be permanently labelled with eYFP. In brachiocephalic trunk sections of 16-week western diet-fed mice, there were less eYFP^+^ Lgals3^+^ macrophages and Ecrg4^+^ fibroblasts localized in the media of the lesions from *Epn1&2-SMC^iDKO^/eYFP^+/-^/ApoE^-/-^*mice compared to that from *eYFP^+/-^/ApoE^-/-^* mice (Figure 2F,G). Meanwhile, significantly more Ecrg4^+^ eYFP^+^ modSMC_myofibroblasts cells were found localized in intima overlying the media in *Epn1&2-SMC^iDKO^/eYFP^+/-^/ApoE^-/-^*mice compared to that of *eYFP^+/-^/ApoE^-/-^* mice (Figure 2G), indicating that modSMC_myofibroblast preferentially localized into the fibrous cap in the atherosclerotic lesions from *Epn1&2- SMC^iDKO^/eYFP^+/-^/ApoE^-/-^*mice. To explore the role of Epsins in the VSMC phenotype switching under atherosclerotic challenges in vitro, aortic VSMCs isolated from *ApoE*^-/-^ mice and tamoxifen injected *Epn1&2-SMC^iDKO^/ApoE^-/-^* mice were treated with 100 μg/mL oxLDL for 72 hrs, followed by western blot. oxLDL treatment led to decreased expression of VSMC contractile makers and such decrease was not as significant in Epsins-deficient VSMCs as that from *ApoE*^-/-^ mice (Figure S5C). Taken together the scRNAseq and biochemical results, we conclude that VSMC-intrinsic Epsins destabilized the contractile phenotypes of VSMCs.

### ModSMC_EC Cells were Functional and Participate in the Repair of Endothelial Injury Caused by Atherosclerosis

Flow cytometry of total cells dissociated from aortae indicated that there were higher proportion of CD31^+^, eYFP^+^ modSMC_EC in total cells from aortae of *Epn1&2- SMC^iDKO^/eYFP^+/-^/ApoE^-/-^*mice compared to that from *eYFP^+/-^/ApoE^-/-^* mice (Figure 3B). Immunostaining of atherosclerotic brachiocephalic trunk sections from *Epn1&2- SMC^iDKO^/eYFP^+/-^/ApoE^-/-^*and *eYFP^+/-^/ApoE^-/-^* mice showed a significantly more CD31^+^ (Figure 3D) or VE-Cadherin^+^ (Figure 3E) cells that were also eYFP^+^ in *Epn1&2-SMC^iDKO^/eYFP^+/-^/ApoE^-/-^* mice compared to that from *eYFP^+/-^/ApoE^-/-^* mice. More importantly, we observed that those CD31, or VE-Cadherin and eYFP double positive cells (modSMC_ECs) were localized in the endothelial layer of blood vessel (Figure 3C-F). Those observations suggested that indeed SMCs can transdifferentiate into endothelial-like cells in atherosclerotic arteries and absence of Epsins promoted such switching. To explore whether Epsins deficiency promotes SMCs to modSMC_EC phenotype switching *in vitro*, we immunostained primary aortic SMCs in long-term culture with αSMA and endothelial markers. Epsins-deficient SMCs showed increased expression of both CD31 and NRP1 compared to wild-type SMCs (Figure S5D). Taken together, those observations suggest that absence of Epsins in SMC promotes the transition of SMC into modSMC_EC and such modSMC_ECs may participate in the repair of EC damages caused by atherosclerotic stimuli.

**Figure 3.**
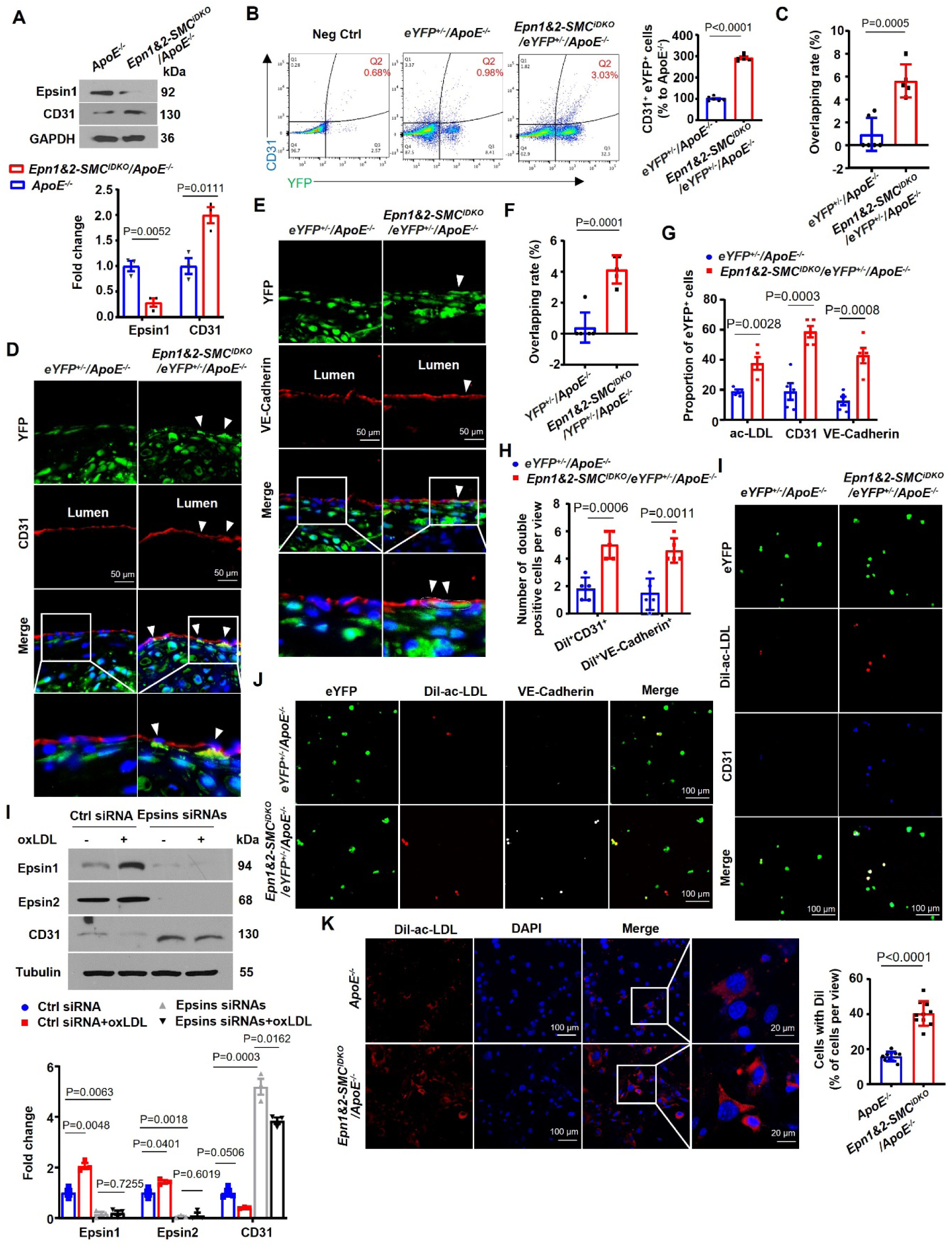
ModSMC_ECs Express Endothelial Markers and are Functional. (A) Immunoblot of Epsin1 and EC marker CD31 on homogenate of aortae from *ApoE^-/-^* and *Epn1&2-SMC^iDKO^/ApoE^-/-^* mice fed a WD for 16 weeks. n=3 mice. (B) Flow cytometry analysis of CD31+, YFP+ cells in aortae of *YFP^+/-^/ApoE^-/-^* and *YFP^+/-^/Epn1&2-SMC^iDKO^/ApoE^-/-^* mice fed on WD for 14 weeks. n=6 mice. (C-F) Immunofluorescence staining of EC markers CD31 (D) and VE-Cadherin (e) on brachiocephalic trunk of *YFP^+/-^/ApoE^-/-^* and *YFP^+/-^/Epn1&2-SMC^iDKO^/ApoE^-/-^*mice fed on WD for 14 weeks. Scale bar=50 μm. n=5-6 mice. White arrowheads indicate YFP^+^ cells that are also positive for CD31 or VE-Cadherin staining. (G-J) Dil-ac-LDL uptake assays, followed by immunofluorescence staining for EC markers CD31 (I) or VE-Cadherin (J) in YFP^+^ cells sorted from total cells dissociated from aortae of *YFP^+/-^/ApoE^-/-^* and *YFP^+/-^/Epn1&2-SMC^iDKO^/ApoE^-/-^* mice fed on WD for 14 weeks. Scale bar=100 μm. (G) Quantification of the proportion of dil^+^, CD31^+^ or VE-Cadherin^+^ cells in YFP^+^ cells as well as (H) the number per view of dil, YFP, CD31 or VE-Cadherin triple positive cells. n=5 mice. (K) Immunoblot of total cell lysate of primary SMC transfected with control or small interference RNAs against Epsin1&2 followed by 100 μg/mL oxLDL stimulation with antibodies indicated. Quantification values were normalized to tubulin expression levels. n=3 independent repeats. (L) Dil-ac-LDL uptake in *in vitro* cultured primary VSMCs from the aortae of *ApoE^-/-^*and *Epn1&2-SMC^iDKO^/ApoE^-/-^* mice. Scale bar=100 and 20 μm, respectively. n=10 independent repeats. SMC, aortic smooth muscle cell; EC, endothelial cell; WD, western diet; Dil-ac-LDL, Dil-acetylated-low density lipoprotein; oxLDL, oxidized low-density lipoprotein. All *P* values were calculated using two-tailed unpaired Student’s *t*-test. Data are mean ± s.d.

One of the fundamental functions of ECs is endocytosis of ac-LDL which helps maintaining homeostasis of blood cholesterol level^58^. Thus, we evaluated whether modSMC_ECs in atherosclerotic plaques can take up ac-LDL. We sorted eYFP^+^ cells from total cells freshly dissociated from the aortae of *eYFP^+/-^/ApoE^-/-^* and *Epn1&2-SMC^iDKO^/eYFP^+/-^ /ApoE^-/-^* mice fed on western diet for 14 weeks. The sorted eYFP^+^ cells were treated with Dil-ac-LDL^59^ for 5 hrs followed by immunostaining with endothelial markers CD31 or VE-Cadherin. There were increased proportion of eYFP^+^ cells that took up ac-LDL, and the proportion of eYFP and CD31 or VE-Cadherin double positive modSMC_ECs were also increased in cells isolated from aortae of *Epn1&2-SMC^iDKO^/eYFP^+/-^/ApoE^-/-^* mice compared to that from *eYFP^+/-^/ApoE^-/-^*. Moreover, there were higher number of Epsins-deficient modSMC_ECs that took up ac-LDL compared to wild-type control modSMC_ECs (Figure 3H- J). Furthermore, each Epsins-deficient modSMC_EC took up more ac-LDL in culture compared to that from wild-type controls (Figure 3K). Western blot revealed that knockdown of Epsins in cultured SMCs lead to increaed expression of CD31 proteins upon oxLDL stimulation (Figure 3L) corroborating earlier findings that Epsins negatively regulate SMC to modSMC_EC transition. Taken together, these data suggest that Epsins deficiency promotes SMCs transdifferentiating to modSMC_ECs and these SMC-originated endothelial-like cells were functional in ingestion of ac-LDL. ModSMC_ECs can integrate into the arterial vessel wall and likely to participate in the repair of atherosclerosis-induced endothelial damage.

### Epsins Suppress the Expression of SMC Markers by Decreasing KLF4 Expression

KLF4 is a critical regulator of SMC phenotypic modulation^7,60–62^. Considering the enrichment of KLF4 in extracellular space evidenced by GeneCards (https://www.genecards.org/cgi-bin/carddisp.pl?gene=KLF4#localization), we performed a Mendelian Randomization analysis to evaluate the relationship between KLF4 and CAD risk (Figure S4A), and observed that higher level of plasma KLF4 protein was causally associated with increased risk of CAD in the general population (*β* = 0.18, *P* = 0.035; Figure 4A). We further observed that KLF4 colocalizes with SMC marker αSMA in atherosclerotic human aortae and the amount of KLF4 protein increased with the advancement of atherosclerosis (Figure 4B). This observation suggests that KLF4 level is increased in SMCs with the progression of atherosclerosis.

**Figure 4.**
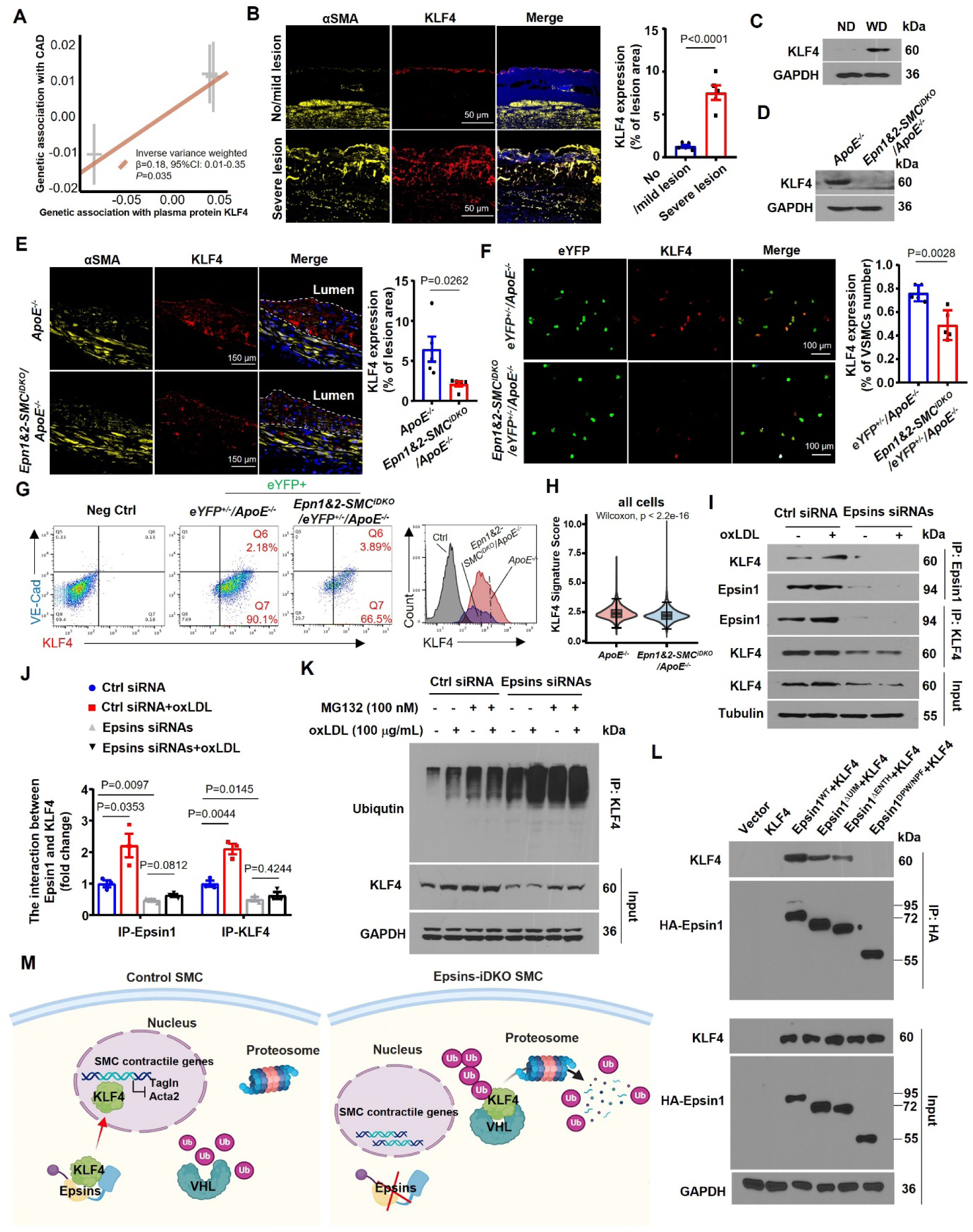
Epsins Stabilizes KLF4 by Interfereing with KLF4 Ubiquitination. (A) Scatter plots for Mendelian Randomization analysis illustrating a putative causal association between plasma KLF4 protein and CAD risk. *P* value was calculated with inverse variance weighted regression using *TwoSampleMR*. (B) Immunofluorescence staining of KLF4 and α-SMA in aortae from patients with no/mild or severe atherosclerotic lesions. Scale bar=50 μm. n=5 samples. (C-D) Immunoblot of KLF4 in the homogenates of aortae from *ApoE^-/-^* mice fed on ND or WD (C) and *ApoE^-/-^* and *Epn1&2-SMC^iDKO^/ApoE^-/-^* mice fed on WD for 16 weeks (D). (E) Immunofluorescence staining for KLF4 in brachiocephalic truncks of SMC-lineage tracing *YFP^+/-^/ApoE^-/-^* and *Epn1&2-SMC^iDKO^/YFP^+/-^/ApoE^-/-^*mice fed on WD for 14 weeks. Scale bar=150 μm. n=5 mice. (F) Immunofluorescence staining of KLF4 in sorted YFP-tagged cells from the aortae of *YFP^+/-^/ApoE^-/-^* and *Epn1&2-SMC^iDKO^/YFP^+/-^/ApoE^-/-^*mice fed on WD for 14 weeks. Scale bar=100 μm. n=5 mice. (G) Flow cytometry plots of VE-Cadherin^+^ and KLF4^+^ cells in cells gated for YFP-postive in total cells dissociated from aortae of *YFP^+/-^/ApoE^- /-^* and *Epn1&2-SMC^iDKO^/YFP^+/-^/ApoE^-/-^*mice fed on WD for 14 weeks. n=6 mice. (H) Differential signature score of KLF4 binding genes between *ApoE^-/-^* and *Epn1&2- SMC^iDKO^/ApoE^-/-^* mice, as revealed by scRNA-seq data. *P* value was calculated by Wilcoxon rank sum test. The signature score was calculated on 1113 target genes with KLF4 binding sites in regulatory regions with *PercentageFeatureSet* function deposited in *Seurat*. (I-J) The interaction between Epsin and KLF4 in primary SMCs from *ApoE^-/-^* and *Epn1&2- SMC^iDKO^/ApoE^-/-^* mice treated with 100 μg/mL oxLDL evaluated with immunoprecipitation followed by western blot. n=3 independent repeats. (K) The KLF4 ubiquitination levels in wild type SMCs transfected with control siRNA or Epsin 1&2 siRNAs following treatment with 100 nM MG132 or 100 μg/mL oxLDL were measured by immunoprecipitation and western blot. (L) HA-tagged Epsin 1 or Epsin 1 domains were co-transfected with pCX4-KLF4 into HEK 293T cells, after 24 hrs, cell lysis was immunoprecipitated with HA antibody, followed by western blot with KLF4 and HA antibodies. (M) Schematic diagram of the proposed mechanism. Epsins stabilize KLF4 and hinder KLF4 ubiquitination by binding to KLF4 through UIM and ENTH domains of Epsin. SMC, aortic smooth muscle cell; EC, endothelial cell; WD, western diet; ND, normal diet; CAD, coronary artery disease; siRNA, small interfering RNA; oxLDL, oxidized low-density lipoprotein. All *P* values were calculated using two-tailed unpaired Student’s *t*-test except (H). Data are mean ± s.d.

Western blot of homogenates of aortae from mice fed on normal or western diet recapitulated the increased expression of KLF4 in atherosclerotic lesions as that observed in human samples (Figure 4C, Figure S6A). Interestingly, such upregulation was almost abrogated in aortae from *Epn1&2-SMC^iDKO^/ApoE^-/-^* mice compared to that from *ApoE^-/-^* mice (Figure 4D,E, Figure S6B). This difference was not due to lower KLF4 mRNA transcript abundance in Epsins-deficient aortae, suggesting a post-transcriptional regulation of KLF4 stability by Epsins (Figure S6C). In addition, we sorted SMCs from aortae of 16-week western diet-fed eYFP-tagged SMC-lineage tracing mice and determined the expression of KLF4 by immunostaining. Both flow cytometric analyses and confocal microscopy showed that about 90.1% of eYFP^+^ SMCs from *eYFP^+/-^/ApoE^-/-^* mice are positive for KLF4. Whereas only 66.5% of eYFP^+^ SMCs from aortae of *Epn1&2-SMC^iDKO^/eYFP^+/-^/ApoE^-/-^* mice were positive for KLF4. Moreover, the KLF4 expression level is lower in eYFP^+^ SMCs from *Epn1&2- SMC^iDKO^/eYFP^+/-^/ApoE^-/-^* mice than that from *eYFP^+/-^/ApoE^-/-^* mice (Figure 4F,G, Figure S6D). We further stimulated SMCs isolated for mouse aortae from mice on normal diet with scrambled or Epsins siRNAs followed by oxLDL. oxLDL treatment induced the upregulation of KLF4 in vitro in SMCs pretreated with scrambled RNA (Figure 4I, Figure S6E). However, both the basal level and oxLDL-induced upregulation of KLF4 were reduced in the absence of Epsins in primary SMC (Figure 4I, Figure S6E). Similarly, immunofluorescence staining showed that KLF4 proteins in primary SMCs were less abundant in SMCs from *Epn1&2- SMC^iDKO^/ApoE^-/-^* mice compared to that from *ApoE^-/-^* mice after oxLDL treatment (Figure S6F). These data indicated that Epsins were crucial for stabilizing the protein level of KLF4 in SMCs. To further explore whether KLF4 is downstream of Epsins in modulating SMC phenotypic modulation, we performed CUT&Tag profiling^30^ against KLF4 in SMCs isolated from aortae of *ApoE^-/-^* and *Epn1&2-SMC^iDKO^/ApoE^-/-^* mice fed on western diet for 16 weeks. After comparing differential binding peaks between the two genotypes, we retrieved 1113 target genes from KLF4 Cut&Tag array and mapped them to scRNAseq dataset as KLF4 binding gene signature (Table S5). Intriguingly, we observed that the KLF4 binding gene signature score was lower in SMC from *Epn1&2-SMC^iDKO^/ApoE^-/-^* mice than that from *ApoE^-/-^* mice (Figure 4H), especially in cell types of SMC, modSMC_EC, modSMC_myofibroblasts, and macrophage cells (Figure S6G). In parallel, we determined the expression pattern of SMC markers in SMCs from *Epn1&2-SMC^iDKO^/ApoE^-/-^*mice after adenovirus-mediated overexpression of KLF4 and observed that SMC markers *Tagln* and *Acta2* were dramatically inhibited upon KLF4 over-expression (Figure S6H). Together, these findings suggest that Epsins promotes SMC phenotype switching in atherosclerosis through increasing KLF4 protein abundance.

### Epsins Binds Directly to KLF4 and Prevent its Ubiquitination and Subsequent Degradation

To explore the molecular mechanisms how Epsins increases the protein abundance of KLF4 in SMCs in atherosclerotic lesions, we performed coimmunoprecipitations from lysates of primary SMCs isolated from *ApoE^-/-^*mice treated with siRNAs to induce the deletion of Epsins followed by oxLDL stimulation. As shown in Figure 4I, we observed a basal binding of KLF4 to Epsin1 in unstimulated SMCs, which was increased in response to oxLDL treatment (Figure 4I,J). KLF4 can be degraded by the ubiquitination-proteosome pathway^63^. To determine whether Epsins control KLF4 stability in SMCs in atherosclerosis through the ubiquitination-proteosome pathway, we transfected SMCs with scrambled and siRNAs against Epsins1&2 followed by treatment with 100 nM proteasome inhibitor MG132 for 6 hrs. Cells were then stimulated with 100 μg/mL oxLDL for 24 hrs. Immunoprecipitation-western showed that Epsins depletion led to decreased KLF4 protein level. oxLDL treatment caused polyubiquitination of KLF4 and such ubiquitination was enhanced upon depletion of Epsins. More importantly, inhibition of proteosome activity increased the protein level of KLF4 in Epsins-depleted SMCs (Figure 4K, Figure S6I,J). Together, those observations suggest that Epsins stabilize KLF4 through inhibiting its ubiquitination and subsequent proteosome- mediated degradation.

We have previously shown that Epsins could recognize ubiquitinated proteins via its ubiquitin-interacting motif (UIM)^21,22^. To determine which Epsins domains are responsible for the interaction with KLF4, we created mammalian expression vectors containing cDNAs encoding HA-tagged full length, ENTH domain, UIM, or ENTH+UIM deletion Epsin1 (HA- Epsin1^WT^, HA-Epsin1^△ENTH^, HA-Epsin1^△UIM^ or HA-Epsin1^DPW/NPF^). We transfected these constructs to HEK 293T cells together with a plasmid expressing KLF4. We performed immunoprecipitation on cell lysates with anti-HA antibody and western blot showed that both ENTH and UIM domain of Epsin1 played a role in the interaction between Epsin1 and KLF4 as the binding between Epsin1^△UIM/△ENTH^ and KLF4 was declined, meanwhile, the binding between Epsin1^DPW/NPF^ and KLF4 was abrogated (Figure 4L, Figure S6K). Taken together, Epsins stabilize KLF4 by binding to KLF4 through its ENTH and UIM domain.

KLF4 ubiquitination is catalyzed by VHL, an E3 ubiquitin ligase, for proteasomal degradation, in breast carcinoma cells^64^. We hypothesize that Epsins inhibits KLF4 ubiquitination by interfering with the interaction between VHL and KLF4. Firstly, scRNAseq data showed that both *Vhl* and *Klf4* expressions in SMC from *Epn1&2-SMC^iDKO^/ApoE^-/-^* mice fed on western diet were significantly increased (Figure S7A). Immunostaining of human aorta sections containing atherosclerotic lesions showed that VHL level in αSMA-positive SMCs correlated strongly with increased disease severity (Figure S7B). To determine whether Epsins interfere with the interaction between VHL and KLF4, we performed co-immunoprecipitation assays in wild-type and Epsins-deficient SMCs using VHL-specific antibody. Epsins deficiency increased VHL protein level in SMCs (Figure S7C,D), and significantly enhanced the interaction between VHL and KLF4 regardless of the presence of oxLDL (Figure S7C,E). Together, those data suggest that loss of Epsins reduced KLF4 expression by interfering with the interaction between VHL and KLF4.

### Epsins Inhibit the Expression of EC Markers in SMCs by Destabilizing OCT4

Given the critical role of OCT4 in controlling the plasticity of SMCs in atherosclerosis^18^, we investigated whether SMC Epsins control OCT4 activity in this process. We have enriched OCT4 binding genes derived from CUT&Tag array (Table S6) and found that OCT4 binding gene signature score in both SMC and modSMC_EC was significantly higher in SMCs from atherosclerotic aortae of *Epn1&2-SMC^iDKO^/ApoE^-/-^* mice than that from *ApoE^-/-^* mice (Figure 5A). Consistently, OCT4 protein level was higher in the homogenate of aorta from *Epn1&2- SMC^iDKO^/ApoE^-/-^* mice compared to that from *ApoE^-/-^* mice, interestingly, we also observed that the RNA level of Oct4 were also upregulated in SMCs isolated from *Epn1&2-SMC^iDKO^/ApoE^-/-^* mice (Figure 5B,C). Furthermore, the increased level of OCT4 protein was observed in aortic arch, thoracic aortae and abdominal aortae from *Epn1&2-SMC^iDKO^/ApoE^-/-^*mice compared with *ApoE^-/-^* mice (Figure 5D). Immunostaining of brachiocephalic trunk from mice fed on western diet for 16 weeks showed that OCT4 was readily detected in the brachiocephalic trunk from *Epn1&2-SMC^iDKO^/ApoE^-/-^* with little detected in brachiocephalic trunk from *ApoE^-/-^* mice and OCT4 co-localized with αSMA (Figure S8A,B). We speculated that Epsins negatively regulate OCT4 protein level regardless of atherosclerotic stimuli. Indeed, the expression of OCT4 in Epsins-knockdown primary SMCs was higher than that in scramble RNA control SMCs regardless of oxLDL treatment (Figure 5E).

**Figure 5.**
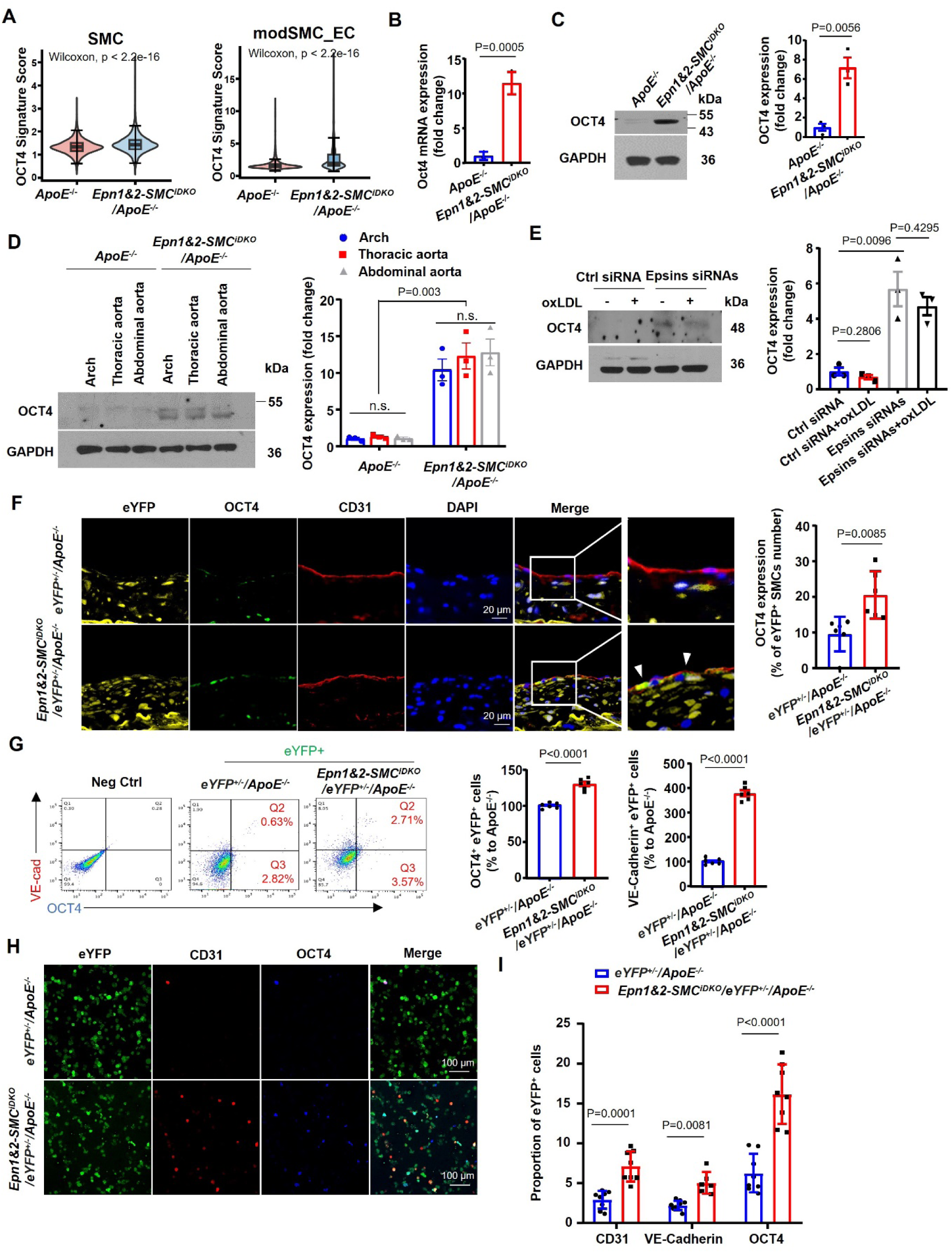
SMC-specific Epsins Deficiencies Promotes Expression of EC Markers in SMCs by Augmenting the OCT4 Expression in Atherosclerotic Plaques. (A) Differential signature score of OCT4 binding genes between *ApoE^-/-^* and *Epn1&2- SMC^iDKO^/ApoE^-/-^* mice across cell types of SMC and modSMC_EC derived from scRNA-seq data. *P* value was calculated by Wilcoxon rank sum test. The signature score was calculated using 898 target genes with OCT4 binding sites in regulatory regions with *PercentageFeatureSet* function deposited in *Seurat*. (B) Relative mRNA level of Oct4 in the cells isolated from the aortae of *ApoE^-/-^* and *Epn1&2-SMC^iDKO^/ApoE^-/-^* mice fed a WD for 16 weeks. n=3 mice with independent repeats. (C-D) Immunoblot analysis of OCT4 expression in either the homogenates of whole aortae (C) or different parts of aortae (D) of *ApoE^-/-^*and *Epn1&2-SMC^iDKO^/ApoE^-/-^*mice fed a WD for 16 weeks. n=3 mice with independent repeats. E, Immunoblot analysis of OCT4 in primary SMC isolated from the aortae of *ApoE^-/-^* and *Epn1&2-SMC^iDKO^/ApoE^-/-^*mice stimulated with or without 100 μg/mL oxLDL for 24 hrs. (B-E) n=3 mice. (F) Immunofluorescence staining of OCT4 in brachiocephalic trunks of YFG- tagged SMC-lineage tracing mice with 14-week WD. Scale bar=20 μm. g, Flow cytometry plots of VE-Cadherin^+^ and Oct4^+^ in YFP^+^ cells in total cells dissociated from aortae of *YFP^+/-^ /ApoE^-/-^* and *Epn1&2-SMC^iDKO^/YFP^+/-^/ApoE^-/-^* mice fed on WD for 14 weeks. (H) Localization of Oct4 and CD31 in sorted YFP^+^ cells from total cells dissociated from aortae of *YFP^+/-^/ApoE^- /-^* and *Epn1&2-SMC^iDKO^/YFP^+/-^/ApoE^-/-^*mice fed on WD for 14 weeks. Scale bar=100 μm. (I) Quantitation of the proportion of CD31*^+^*, VE-Cadherin*^+^* and OCT4*^+^* in YFP*^+^* cells. n=6-8 mice. (F-H) n=6 mice. SMC, aortic smooth muscle cell; EC, endothelial cell; WD, western diet; oxLDL, oxidized low-density lipoprotein. All *P* values were calculated using two-tailed unpaired Student’s *t*-test except (A). Data are mean ± s.d.

To explore the role of OCT4 in SMC phenotype modulation, we immunostained brachiocephalic trunk sections from *eYFP^+/-^/ApoE^-/-^* and *Epn1&2-SMC^iDKO^/eYFP^+/-^/ApoE^-/-^* mice with anti-CD31 and OCT4 antibodies. OCT4 expressed highly in eYFP^+^ CD31^+^ cells localized in the intima of plaque of brachiocephalic trunk from *Epns-SMC^iDKO^/eYFP^+/-^/ApoE^-/-^* mice (Figure 5F). To corroborate those immunostaining findings, we dissociated aortae cells and stained with anti-VE-cadherin and observed that about 2.71% of eYFP^+^ SMCs were OCT4 and VE-cadherin double positive in *Epn1&2-SMC^iDKO^/eYFP^+/-^/ApoE^-/-^*mice, whereas only 0.63% were found in brachiocephalic trunk from *eYFP^+/-^/ApoE^-/-^* mice (Figure 5G). In addition, we sorted eYFP-positive cells from SMC-lineage tracing mice fed on western diet for 16 weeks and stained with CD31/VE-Cadherin and OCT4. There were more OCT4^+^ CD31^+^ and OCT4^+^ VE-Caherin^+^ eYFP-tagged cells from *Epn1&2-SMC^iDKO^/eYFP^+/-^/ApoE^-/-^*than that from *eYFP^+/-^/ApoE^-/-^* mice (Figure 5H,I Figure S8C). Taken together, OCT4 preferentially localized in EC marker-positive modSMC_ECs suggesting that OCT4 may play a role in SMC to modSMC_EC modulation.

To further explore whether OCT4 is downstream of Epsins to suppress the expression of SMC contractile markers as well as SMC to endothelial transdifferentiation, we performed both qRT-PCR and western blot in SMCs isolated from *Epn1&2-SMC^iDKO^/ApoE^-/-^* mice treated with tamoxifen and OCT4 siRNA. Knocking down of *Oct4* in Epsins-deficient SMC led to decrease of SMC contractile markers as well as EC marker CD31 (Figure S8D,E).

### SMC-Specific Epsins Deficiency Reduced the Size of Atherosclerotic Lesion and Increased the Plaque Stability in *vivo*

Given our observation that Epsins expression was upregulated in SMC in atherosclerotic lesions and SMC-specific Epsins deficiency selectively increase SMC transition to endothelial-like cells which in turn participate in the repair of endothelial injury caused by atherogenic stimuli, we speculated that Epsins deficiency in SMC would improve the outcome of experimental atherosclerosis *in vivo*. To test this, *ApoE^-/-^* mice and tamoxifen injected *Epn1&2- SMC^iDKO^/ApoE^-/-^* mice were fed a western diet for 9, 16, 20 weeks. Assessment of the *en face* lesion area in whole aorta revealed that atherosclerotic lesion was significantly smaller in *Epn1&2-SMC^iDKO^/ApoE^-/-^* mice compared to *ApoE^-/-^*mice (*P* = 0.0012 with 9-week western diet, *P* = 0.0247 with 16-week western diet, *P* = 0.0003 with 20-week western diet, compared with control *ApoE^-/-^* mice; Figure 6A). Furthermore, examination of aortic root lesions demonstrated that tamoxifen injected *Epn1&2-SMC^iDKO^/ApoE^-/-^* mice had a 56.57% reduction in lesion area in comparison to *ApoE^-/-^* mice (Figure 6B,C). Using Oil Red O staining, we also observed significant reduction in lipid loading in the lesion of sinus and brachiocephalic trunk in *Epn1&2-SMC^iDKO^/ApoE^-/-^* mice compared to *ApoE^-/-^* mice (Figure 6D,E).

**Figure 6.**
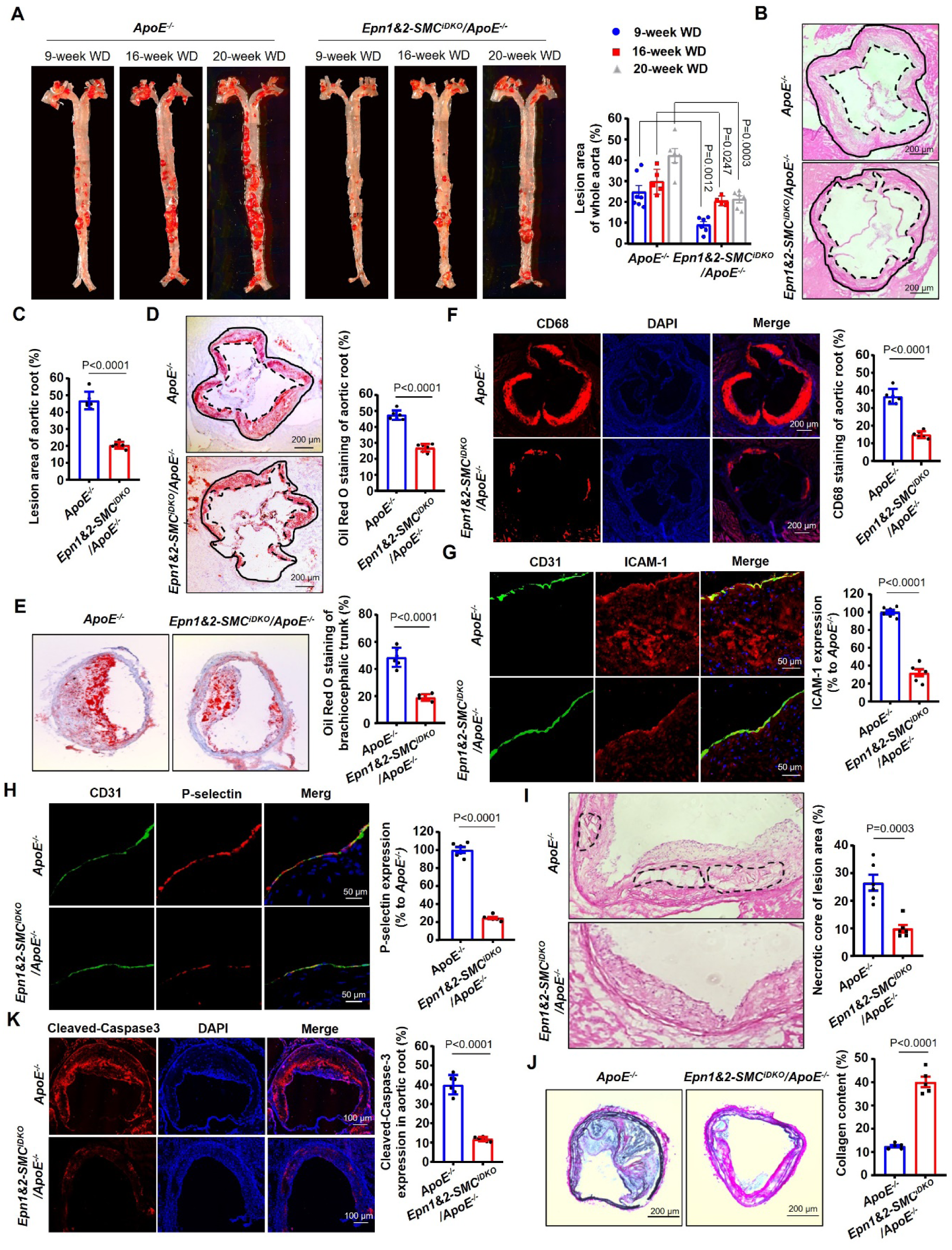
SMC-specific Epsin1&2 Deficiency Reduces Atherosclerotic Plaques and Enhances the Stability of Lesions in Mice. *ApoE^-/-^* and *Epn1&2-SMC^iDKO^/ApoE^-/-^*mice were fed a WD for 9, 16 and 20 weeks. The sections of aortic root and brachiocephalic trunks were collected from the *ApoE^-/-^* and *Epn1&2- SMC^iDKO^/ApoE^-/-^* mice fed a WD for 16 weeks. (A) *En face* Oil Red O staining of the whole aortae was calculated. (A)The sections of aortic root and brachiocephalic trunks were collected stained en face with Oil Red O. (B-C) Hematoxylin and eosin staining of aortic roots showed the size of atherosclerotic lesion. (D-E) Oil Red O staining of aortic roots (D) and brachiocephalic trunks (E) were used to show the lipid accumulation in the lesionof aortae of *ApoE^-/-^* and *Epn1&2-SMC^iDKO^/ApoE^-/-^* mice fed on WD for 16 weeks. (F) Immunofluorescence staining of CD68 were performed to show the inflammation in the lesion of aortic root from aortae of *ApoE^-/-^* and *Epn1&2-SMC^iDKO^/ApoE^-/-^*mice fed on WD for 16 weeks. (G-H) Immunofluorescence staining for CD31 and ICAM-1 (G) or P-selectin (H) in dissected aortic roots from aortae of *ApoE^-/-^* and *Epn1&2-SMC^iDKO^/ApoE^-/-^* mice fed on WD for 16 weeks. Scale bar=50 μm. (I) Hematoxylin and eosin staining of aortic roots showed the necrotic cores in the lesion. Necrotic cores were outlined in black dash. (J) Verhoeff-Van Gieson’s staining of brachiocephalic trunks was performed to show the stability of the lesion of aortae from *ApoE^-/-^*and *Epn1&2-SMC^iDKO^/ApoE^-/-^* mice fed on WD for 16 weeks. Scale bar=200 mm. (K) Immunofluorescence staining of cleaved (active)-caspase3 in aortic root from *ApoE^-/-^* and *Epn1&2-SMC^iDKO^/ApoE^-/-^*mice fed on WD for 16 weeks. Scale bar=100 μm. SMC, aortic smooth muscle cell; EC, endothelial cell; WD, western diet. All *P* values were calculated using two-tailed unpaired Student’s *t*-test. Data are mean ± s.d. n=6 mice.

To evaluate the inflammatory profile of the atherosclerotic lesions of these mice, we assessed lesion composition by immunostaining with markers of macrophages (CD68) within aortic sinus lesions. Consistent with a reduction in atherosclerotic progression, we detected a reduction in macrophage area by 59.2% in tamoxifen injected *Epn1&2-SMC^iDKO^/ApoE^-/-^* mice (*P* < 0.0001 vs *ApoE^-/-^*mice; Figure 6F). Last, we observed a reduction in ICAM-1 and P-selectin staining in the endothelial layer of aortic root lesions of tamoxifen injected *Epn1&2- SMC^iDKO^/ApoE^-/-^* mice compared to *ApoE^-/-^* mice (Figure 6G-H), which was consistent with the reduced recruitment and accumulation of macrophages in the atherosclerotic plaques of these mice. Together, these data demonstrated that deficiency of Epsin1&2 in SMC could protect against atherosclerotic progression induced by a western diet.

In addition to reduction of lesion size, a thick fibrous cap as well as a smaller necrotic core are important features of stable plaques that are less likely to be disrupted to cause thrombosis^65^. Our scRNAseq results showed that loss of *Epsin1&2* in SMCs increased the SMC-derived myofibroblasts which is beneficial to the protective fibrous cap formation^12^ (Figure 1D, Figure S2D). Deficiency of Epsins in SMC also led to increased thickness of the fibrous cap of atherosclerotic lesions in *Epn1&2-SMC^iDKO^/ApoE^-/-^* mice compared to that in *ApoE^-/-^* mice fed on western diet (Figure 2E) To assess plaque stability, we analyzed the size of necrotic core, plaque collagen content, accumulation of contractile SMC content and dead cell content. Epsins specific knockout in SMCs: 1) significantly reduced the necrotic core area to total plaque area ratio as determined by hematoxylin and eosin staining (Figure 6I), 2) markedly elevated the total plaque collagen content as determined by Van Gieson’s staining (Figure 6J), and 3) significantly decreased the dead cell content which was marked by cleaved-Caspase3 (Figure 6K) in atherosclerotic lesions.

## Discussion

Our previous studies have elucidated an atheroprone function of Epsins in both ECs^20,21^, and macrophages^22,66^ in the pathogenesis of atherosclerosis. However, as the major source of plaque cells and extracellular matrix (ECM) at all stages of atherosclerosis, the role of SMC-intrinsic Epsins in pathogenesis of atherosclerosis remains largely unknown. By integrating scRNAseq data with GWAS, we discovered that Epsins-deficiency specifically in VSMCs led to suppressed expression of genes associated with increased CAD risks, highlighting the pivotal role of VSMC-intrinsic Epsins in the pathological process leading to CAD. Using single-cell genomics and mouse strains with VSMC-specific Epsins deficiency and lineage tracing, we revealed VSMCs can transdifferentiate into endothelial-like cells. Specifically, loss of Epsins results in a higher proportion of SMC-derived endothelial-like cells and myofibroblasts, particularly in advanced atherosclerotic lesions. While the SMC-derived endothelial-like cells may participate in the repair of endothelial damage, the myofibroblasts transdifferentiated from SMCs stabilizes atherosclerotitc plaque cap. Both cell types are athero-protective. Therefore, the SMC-intrinsic Epsins promote the pathogenesis of atherosclerosis at least in part through inhibition of phenotype switching of SMCs into athero-protective cell types.

Using single-cell genomics and mouse strains with SMC-specific Epsins deficiency and lineage tracing, we revealed SMCs can transdifferentiate into endothelial-like cells. Specifically, loss of Epsins results in a higher proportion of SMC-derived endothelial-like cells and myofibroblasts, particularly in advanced atherosclerotic lesions. Previous studies reported SMC-derived *Vcam*^+^ cells, which were assumed to be SEMs (<u>s</u>tem cell, <u>e</u>ndothelial cell and <u>m</u>onocytes/macrophage differentiation, termed de-differentiated SMCs) defined as an intermediate cell state, in human atherosclerotic plaques and mouse atherosclerotic models^13^. Of greatest significance, through a series of trajectory analyses of our scRNAseq data, we show that the absence of Epsins not only increase the transition from SMC to modSMC_EC, but also stabilized SMC-derived endothelial-like cell population. Harnessing a lineage tracing mice, in combination with flow cytometry cell sorting and endothelial functional assessment, as well as confocal microscopy of aortae sections, we revealed that SMC-derived endothelial-like cells showed basic EC functions and may integrate into vascular vessel wall to participate the reparation of endothelial injury caused by atherosclerotic stimuli. Taken together, our studies suggest that the molecular features of SMC-derived endothelial-like cells represent a unique transitional state from SMCs to endothelial-like cells in the milieu of atherosclerosis. Epsins’ deficiency enhanced the conversion of SMCs into such SMC-derived endothelial-like cells. Anatomically, there are multiple layers of SMCs whereas there is only one single layer of endothelial cell in arterial wall. This is reflected by the scRNAseq data that SMC is the largest population while ECs is a minor one. Despite the fact that the proportion of modSMC_EC is low among the whole modulated SMC-derived cell population, those modSMC_ECs are located in the vicinity of damaged endothelial cells. Therefore, those small number of modSMC_ECs play a pivotal role in the repair of injured arterial endothelial wall. The mechanisms why modSMC_ECs are mostly located near arterial injury sites deserves further investigation.

In the current study, Mendelian Randomization analysis showed that the elevated plasma KLF4 protein constitute as a risk factor of CAD in the general population. Consistently, KLF4 in SMCs was abundant in both human and mouse atherosclerotic lesions which correlated with the upregulation of Epsins. More importantly, the upregulation of KLF4 in response to atherogenic stimuli was attenuated in the absence of Epsins. Given Epsins’ classical role as a membrane-associated endocytic adaptor, they are unlikely to regulate *KLF4* at the transcriptional level. It has been reported previously that once KLF4 is expelled from the nuclear, it is quickly ubiquitinated followed by proteosome-mediated degradation in mouse blastocysts^67^. In the absence of Epsins, both KLF4 expression level and nuclei localization were reduced in oxLDL treated SMCs, suggesting that Epsins are critical for maintaining KLF4 protein level. In contractile SMCs, KLF4 protein level is kept in check by VHL ubiquitin E3 ligase which serve to prevent the conversion of those cells into synthetic phenotypes^68^. Our current data supports a model in which Epsins interacts constitutively with KLF4 via its UIM and ENTH domains, interfering with KLF4-VHL interaction, thus reducing KLF4 ubiquitination and degradation.

KLF4 as a transcription factor has been implicated in SMC phenotype modulation^7,69^. In our study, we showed that phenotypically modulated SMCs transdifferentiate to multiple phenotypes, including cells that express markers of myofibroblast, macrophage, and EC in a mouse model of atherosclerosis. Forced ectopic expression of KLF4 in Epsins-deficient SMCs induced a marked reduction in SMCs contractile phenotype, while it had no effect on endothelial-like SMC phenotype. However, the role of KLF4 in regulation of gene expression and coordination with other transcriptions factors is context-dependent^60,70,71^. In addition to transcriptional regulation, KLF4 functions as a scaffolding protein to recruit other transcriptional regulators to promoters of SMC marker genes in response to different environmental cues^72^. KLF4/OCT4 complex is sufficient and necessary to generate induced pluripotent stem cells from several cell types, such as dermal papilla cells and adult neural stem cells^73, 74,75^. However, previous studies have shown that loss of OCT4 within SMC had virtually completely opposite overall effects on lesion pathogenesis as compared to SMC specific loss of KLF4^15^. In our current study, knockdown of OCT4 in SMCs resulted in reduced SMC markers and EC marker expression. Our data supported that KLF4 and OCT4 controls opposite aspects of SMC phenotypic transitions in the atherosclerotic context^7,18^. It remains unclear how KLF4 and OCT4 coordinate to control the transdifferentiaton of VSMCs. It has been shown that KLF4 can bind to the promoter region of OCT4 and enhance OCT4 transcription^18^. Nevertheless, we found that SMC Epsins deficiency led to decreased KLF4 but increased OCT4 protein level in SMC-derived endothelial-like cells. Though OCT4 expression was hardly detectable in the aorta of the *ApoE^-/-^* control mice which is in line with earlier findings that Oct4 is barely detectable in most adult mouse organs^76^, nevertheless, OCT4 protein level was increased dramatically in aortae from Epsins-deficient mice. The underlying molecular mechanisms by which Epsins control the protein level of Oct4 merits further investigation in relation to KLF4 level. In addition, future studies are warranted to comprehensively evaluate the predictive value of the therapeutic effects of targeting VSMC Epsins with siRNA nanoparticles to halt atherosclerosis progression.

In this study, we discovered that the expression level of Epsins in SMCs tightly correlated with the severity of the disease in both human and mouse atherosclerotic aortae. Notably, selective loss of *Epsin1 and 2* within SMCs led to 1) reduced lesion size and lipid load, 2) enhanced stability of the plaque, including increased fibrous cap thickness and decreased necrotic core, 3) reduced proinflammatory macrophage and the number of dead cells in the necrotic core, 4) increased *Myh11^+^* and *Acta2*^+^ cells within the fibrous cap. Taken together our discovery that Epsins differentially control the protein level of KLF4 and OCT4 in aortic SMCs, and previously established disparate role of Klf4 and Oct4 in the pathogenesis of atherosclerosis^7,18,19^ support our findings, an atheroprone function of SMC Epsins.

In summary, our study reveals a novel cell state during SMC phenotypic switching and identifies potential therapeutic targets to repair the dysfunctional endothelium in atherosclerosis. Epsins are critical for SMC dedifferentiation in atherosclerosis disease progression by protecting KLF4 from ubiquitination and proteasomal degradation. The absence of Epsins in SMCs resulted in the loss of KLF4 and hampered the progression of atherosclerosis in *Epn1&2-SMC^iDKO^/ApoE^-/-^* mice. These insights may pave the way for targeted SMC-Epsins inhibition as a novel therapeutic treatment of CAD.

## Acknowledgments

We thank the Flow Cytometry Core at Boston Children’s Hospital for the use of the LSRII, BGI Hong Kong for the scRNA-sequencing, and the Biopolymers Facility at Harvard Medical School for quality control analysis of RNA samples. Dr. John Shyy from University of California, San Diego provided KLF4 AAV construct. Dr. Gary K. Owens from University of Virginia provided Rosa26Stop-floxed eYFP reporter stain of mice. Xinlei Gao and Kaifu Chen from Boston Children’s Hospital/Harvard Medical School provided bioinformatic suggestions.

## Author Contributions

B.W. and H.C. conceived and designed the experiments. B.W., K.C. and B.Z. performed most of the experiments. B.W. primarily contributed to the in vivo molecular mechanism. B.W. and K.C. primarily contributed to in vitro data in atherosclerosis analysis. B.Z. prepared samples for the scRNA-sequencing and did cDNA library construction. M.D. analyzed the scRNA-seq data and performed bioinformatic work. H.W., K.L. and B.S. helped with immunostaining. Y.S. helped with the primary SMC isolation. B.W. and D.W. analyzed the data and provided comments. H.W., and Y.D. worked on part of the molecular mechanism investigation. K.L. and S.W. performed the mouse genotyping and colony maintenance. B.W., D.W., M.D., D.B.C., Y.C. and H.C. wrote and edited the article. All the authors reviewed and provided feedback on the article.

## Supplemental Material

Document S1. Figures S1–S9 and Tables S1 and S2

Table S3. Excel file containing additional data too large to fit in a PDF, related to Figure 2

Table S4. Excel file containing even more data too large to fit in a PDF, related to Figure 2

Table S5. Excel file containing even more data too large to fit in a PDF, related to Figure 2

Table S6. Excel file containing even more data too large to fit in a PDF, related to Figure 5

---

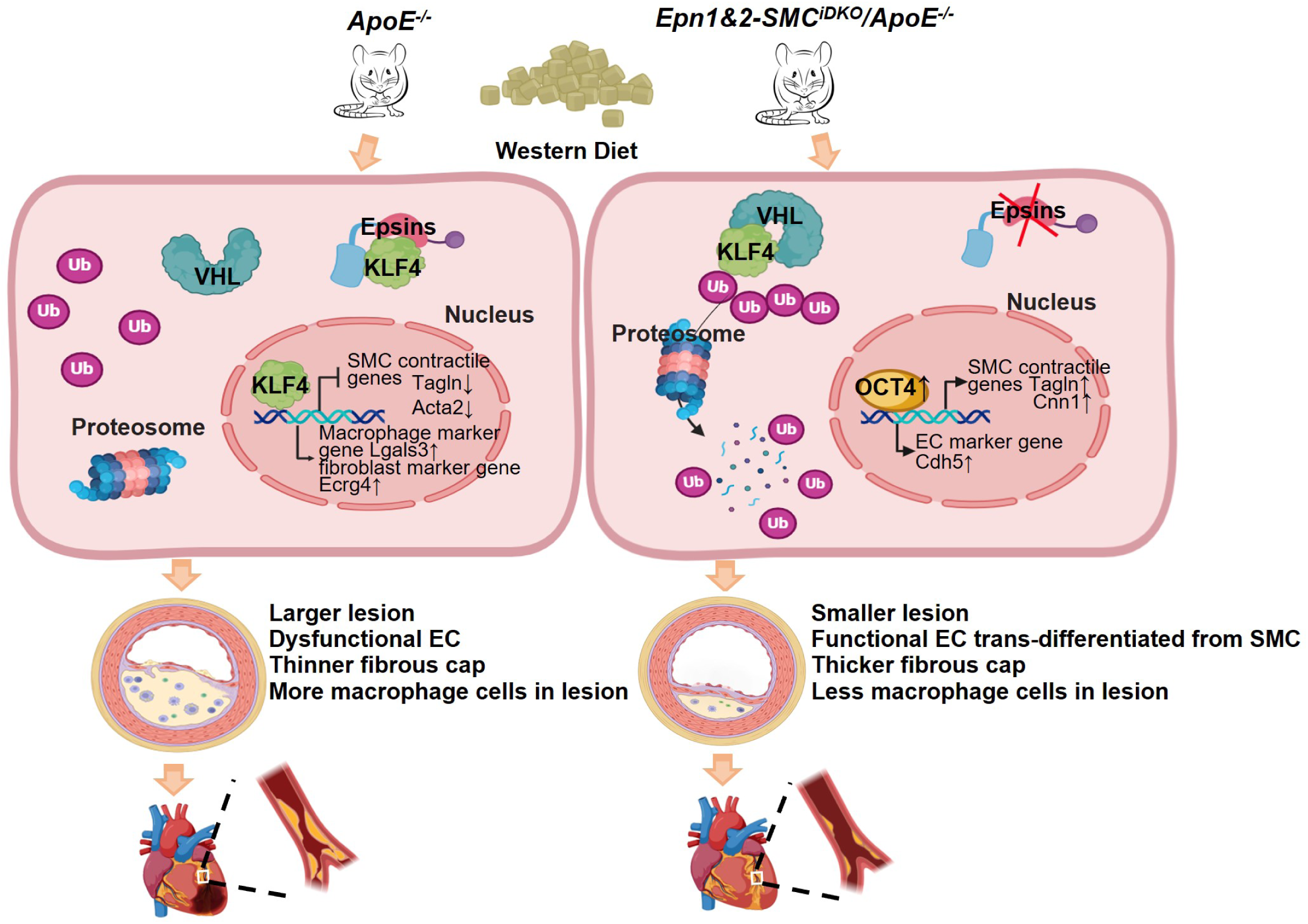

## References

1. Frak, W., Wojtasinska, A., Lisinska, W., Mlynarska, E., Franczyk, B., and Rysz, J. (2022). Pathophysiology of Cardiovascular Diseases: New Insights into Molecular Mechanisms of Atherosclerosis, Arterial Hypertension, and Coronary Artery Disease. Biomedicines 10. 10.3390/biomedicines10081938.

2. Kong, P., Cui, Z.Y., Huang, X.F., Zhang, D.D., Guo, R.J., and Han, M. (2022). Inflammation and atherosclerosis: signaling pathways and therapeutic intervention. Signal Transduct Target Ther 7, 131. 10.1038/s41392-022-00955-7.

3. Badimon, L., and Vilahur, G. (2014). Thrombosis formation on atherosclerotic lesions and plaque rupture. J Intern Med 276, 618–632. 10.1111/joim.12296.

4. Kowara, M., and Cudnoch-Jedrzejewska, A. (2021). Pathophysiology of Atherosclerotic Plaque Development-Contemporary Experience and New Directions in Research. Int J Mol Sci 22. 10.3390/ijms22073513.

5. Chappell, J., Harman, J.L., Narasimhan, V.M., Yu, H., Foote, K., Simons, B.D., Bennett, M.R., and Jorgensen, H.F. (2016). Extensive Proliferation of a Subset of Differentiated, yet Plastic, Medial Vascular Smooth Muscle Cells Contributes to Neointimal Formation in Mouse Injury and Atherosclerosis Models. Circ Res 119, 1313–1323. 10.1161/CIRCRESAHA.116.309799.

6. Allahverdian, S., Chaabane, C., Boukais, K., Francis, G.A., and Bochaton-Piallat, M.L. (2018). Smooth muscle cell fate and plasticity in atherosclerosis. Cardiovasc Res 114, 540–550. 10.1093/cvr/cvy022.

7. Shankman, L.S., Gomez, D., Cherepanova, O.A., Salmon, M., Alencar, G.F., Haskins, R.M., Swiatlowska, P., Newman, A.A., Greene, E.S., Straub, A.C., et al. (2015). KLF4- dependent phenotypic modulation of smooth muscle cells has a key role in atherosclerotic plaque pathogenesis. Nat Med 21, 628–637. 10.1038/nm.3866.

8. Gomez, D., and Owens, G.K. (2012). Smooth muscle cell phenotypic switching in atherosclerosis. Cardiovasc Res 95, 156–164. 10.1093/cvr/cvs115.

9. Rong, J.X., Shapiro, M., Trogan, E., and Fisher, E.A. (2003). Transdifferentiation of mouse aortic smooth muscle cells to a macrophage-like state after cholesterol loading. Proc Natl Acad Sci U S A 100, 13531–13536. 10.1073/pnas.1735526100.

10. Pidkovka, N.A., Cherepanova, O.A., Yoshida, T., Alexander, M.R., Deaton, R.A., Thomas, J.A., Leitinger, N., and Owens, G.K. (2007). Oxidized phospholipids induce phenotypic switching of vascular smooth muscle cells in vivo and in vitro. Circ Res 101, 792–801. 10.1161/CIRCRESAHA.107.152736.

11. Bennett, M.R., Sinha, S., and Owens, G.K. (2016). Vascular Smooth Muscle Cells in Atherosclerosis. Circ Res 118, 692–702. 10.1161/CIRCRESAHA.115.306361.

12. Wirka, R.C., Wagh, D., Paik, D.T., Pjanic, M., Nguyen, T., Miller, C.L., Kundu, R., Nagao, M., Coller, J., Koyano, T.K., et al. (2019). Atheroprotective roles of smooth muscle cell phenotypic modulation and the TCF21 disease gene as revealed by single-cell analysis. Nat Med 25, 1280–1289. 10.1038/s41591-019-0512-5.

13. Pan, H., Xue, C., Auerbach, B.J., Fan, J., Bashore, A.C., Cui, J., Yang, D.Y., Trignano, S.B., Liu, W., Shi, J., et al. (2020). Single-Cell Genomics Reveals a Novel Cell State During Smooth Muscle Cell Phenotypic Switching and Potential Therapeutic Targets for Atherosclerosis in Mouse and Human. Circulation 142, 2060–2075. 10.1161/CIRCULATIONAHA.120.048378.

14. Tetreault, M.P., Yang, Y., and Katz, J.P. (2013). Kruppel-like factors in cancer. Nat Rev Cancer 13, 701–713. 10.1038/nrc3582.

15. Alencar, G.F., Owsiany, K.M., Karnewar, S., Sukhavasi, K., Mocci, G., Nguyen, A.T., Williams, C.M., Shamsuzzaman, S., Mokry, M., Henderson, C.A., et al. (2020). Stem Cell Pluripotency Genes Klf4 and Oct4 Regulate Complex SMC Phenotypic Changes Critical in Late-Stage Atherosclerotic Lesion Pathogenesis. Circulation 142, 2045–2059. 10.1161/CIRCULATIONAHA.120.046672.

16. Salmon, M., Gomez, D., Greene, E., Shankman, L., and Owens, G.K. (2012). Cooperative binding of KLF4, pELK-1, and HDAC2 to a G/C repressor element in the SM22alpha promoter mediates transcriptional silencing during SMC phenotypic switching in vivo. Circ Res 111, 685-696. 10.1161/CIRCRESAHA.112.269811.

17. Hu, D., Gur, M., Zhou, Z., Gamper, A., Hung, M.C., Fujita, N., Lan, L., Bahar, I., and Wan, Y. (2015). Interplay between arginine methylation and ubiquitylation regulates KLF4-mediated genome stability and carcinogenesis. Nat Commun 6, 8419. 10.1038/ncomms9419.

18. Cherepanova, O.A., Gomez, D., Shankman, L.S., Swiatlowska, P., Williams, J., Sarmento, O.F., Alencar, G.F., Hess, D.L., Bevard, M.H., Greene, E.S., et al. (2016). Activation of the pluripotency factor OCT4 in smooth muscle cells is atheroprotective. Nat Med 22, 657–665. 10.1038/nm.4109.

19. Shin, J., Tkachenko, S., Gomez, D., Tripathi, R., Owens, G.K., and Cherepanova, O.A. (2023). Smooth muscle cells-specific loss of OCT4 accelerates neointima formation after acute vascular injury. Front Cardiovasc Med 10, 1276945. 10.3389/fcvm.2023.1276945.

20. Dong, Y., Wang, B., Du, M., Zhu, B., Cui, K., Li, K., Yuan, K., Cowan, D.B., Bhattacharjee, S., Wong, S., et al. (2023). Targeting Epsins to Inhibit Fibroblast Growth Factor Signaling While Potentiating Transforming Growth Factor-beta Signaling Constrains Endothelial-to-Mesenchymal Transition in Atherosclerosis. Circulation 147, 669–685. 10.1161/CIRCULATIONAHA.122.063075.

21. Dong, Y., Lee, Y., Cui, K., He, M., Wang, B., Bhattacharjee, S., Zhu, B., Yago, T., Zhang, K., Deng, L., et al. (2020). Epsin-mediated degradation of IP3R1 fuels atherosclerosis. Nat Commun 11, 3984. 10.1038/s41467-020-17848-4.

22. Cui, K., Gao, X., Wang, B., Wu, H., Arulsamy, K., Dong, Y., Xiao, Y., Jiang, X., Malovichko, M.V., Li, K., et al. (2023). Epsin Nanotherapy Regulates Cholesterol Transport to Fortify Atheroma Regression. Circ Res 132, e22–e42. 10.1161/CIRCRESAHA.122.321723.

23. Kawai-Kowase, K., Ohshima, T., Matsui, H., Tanaka, T., Shimizu, T., Iso, T., Arai, M., Owens, G.K., and Kurabayashi, M. (2009). PIAS1 mediates TGFbeta-induced SM alpha-actin gene expression through inhibition of KLF4 function-expression by protein sumoylation. Arterioscler Thromb Vasc Biol 29, 99–106. 10.1161/ATVBAHA.108.172700.

24. Adam, P.J., Regan, C.P., Hautmann, M.B., and Owens, G.K. (2000). Positive- and negative-acting Kruppel-like transcription factors bind a transforming growth factor beta control element required for expression of the smooth muscle cell differentiation marker SM22alpha in vivo. J Biol Chem 275, 37798–37806. 10.1074/jbc.M006323200.

25. Wirth, A., Benyo, Z., Lukasova, M., Leutgeb, B., Wettschureck, N., Gorbey, S., Orsy, P., Horvath, B., Maser-Gluth, C., Greiner, E., et al. (2008). G12-G13-LARG-mediated signaling in vascular smooth muscle is required for salt-induced hypertension. Nat Med 14, 64–68. 10.1038/nm1666.

26. Deaton, R.A., Bulut, G., Serbulea, V., Salamon, A., Shankman, L.S., Nguyen, A.T., and Owens, G.K. (2023). A New Autosomal Myh11-CreER(T2) Smooth Muscle Cell Lineage Tracing and Gene Knockout Mouse Model-Brief Report. Arterioscler Thromb Vasc Biol 43, 203–211. 10.1161/ATVBAHA.122.318160.

27. Li, Z., Martin, M., Zhang, J., Huang, H.Y., Bai, L., Zhang, J., Kang, J., He, M., Li, J., Maurya, M.R., et al. (2017). Kruppel-Like Factor 4 Regulation of Cholesterol-25- Hydroxylase and Liver X Receptor Mitigates Atherosclerosis Susceptibility. Circulation 136, 1315–1330. 10.1161/CIRCULATIONAHA.117.027462.

28. Stuart, T., Butler, A., Hoffman, P., Hafemeister, C., Papalexi, E., Mauck, W.M., 3rd, Hao, Y., Stoeckius, M., Smibert, P., and Satija, R. (2019). Comprehensive Integration of Single-Cell Data. Cell 177, 1888-1902 e1821. 10.1016/j.cell.2019.05.031.

29. Korsunsky, I., Millard, N., Fan, J., Slowikowski, K., Zhang, F., Wei, K., Baglaenko, Y., Brenner, M., Loh, P.R., and Raychaudhuri, S. (2019). Fast, sensitive and accurate integration of single-cell data with Harmony. Nat Methods 16, 1289–1296. 10.1038/s41592-019-0619-0.

30. Kaya-Okur, H.S., Wu, S.J., Codomo, C.A., Pledger, E.S., Bryson, T.D., Henikoff, J.G., Ahmad, K., and Henikoff, S. (2019). CUT&Tag for efficient epigenomic profiling of small samples and single cells. Nat Commun 10, 1930. 10.1038/s41467-019-09982-5.

31. 31. La Manno, G., Soldatov, R., Zeisel, A., Braun, E., Hochgerner, H., Petukhov, V., Lidschreiber, K., Kastriti, M.E., Lonnerberg, P., Furlan, A., et al. (2018). RNA velocity of single cells. Nature 560, 494–498. 10.1038/s41586-018-0414-6.

32. Bergen, V., Lange, M., Peidli, S., Wolf, F.A., and Theis, F.J. (2020). Generalizing RNA velocity to transient cell states through dynamical modeling. Nat Biotechnol 38, 1408–1414. 10.1038/s41587-020-0591-3.

33. Gulati, G.S., Sikandar, S.S., Wesche, D.J., Manjunath, A., Bharadwaj, A., Berger, M.J., Ilagan, F., Kuo, A.H., Hsieh, R.W., Cai, S., et al. (2020). Single-cell transcriptional diversity is a hallmark of developmental potential. Science 367, 405–411. 10.1126/science.aax0249.

34. Qiu, X., Mao, Q., Tang, Y., Wang, L., Chawla, R., Pliner, H.A., and Trapnell, C. (2017). Reversed graph embedding resolves complex single-cell trajectories. Nat Methods 14, 979–982. 10.1038/nmeth.4402.

35. Jin, S., Guerrero-Juarez, C.F., Zhang, L., Chang, I., Ramos, R., Kuan, C.H., Myung, P., Plikus, M.V., and Nie, Q. (2021). Inference and analysis of cell-cell communication using CellChat. Nat Commun 12, 1088. 10.1038/s41467-021-21246-9.

36. Nelson, C.P., Goel, A., Butterworth, A.S., Kanoni, S., Webb, T.R., Marouli, E., Zeng, L., Ntalla, I., Lai, F.Y., Hopewell, J.C., et al. (2017). Association analyses based on false discovery rate implicate new loci for coronary artery disease. Nat Genet 49, 1385–1391. 10.1038/ng.3913.

37. Nikpay, M., Goel, A., Won, H.H., Hall, L.M., Willenborg, C., Kanoni, S., Saleheen, D., Kyriakou, T., Nelson, C.P., Hopewell, J.C., et al. (2015). A comprehensive 1,000 Genomes-based genome-wide association meta-analysis of coronary artery disease. Nat Genet 47, 1121–1130. 10.1038/ng.3396.

38. Myocardial Infarction, G., Investigators, C.A.E.C., Stitziel, N.O., Stirrups, K.E., Masca, N.G., Erdmann, J., Ferrario, P.G., Konig, I.R., Weeke, P.E., Webb, T.R., et al. (2016). Coding Variation in ANGPTL4, LPL, and SVEP1 and the Risk of Coronary Disease. N Engl J Med 374, 1134-1144. 10.1056/NEJMoa1507652.

39. de Leeuw, C.A., Mooij, J.M., Heskes, T., and Posthuma, D. (2015). MAGMA: generalized gene-set analysis of GWAS data. PLoS Comput Biol 11, e1004219. 10.1371/journal.pcbi.1004219.

40. Ferkingstad, E., Sulem, P., Atlason, B.A., Sveinbjornsson, G., Magnusson, M.I., Styrmisdottir, E.L., Gunnarsdottir, K., Helgason, A., Oddsson, A., Halldorsson, B.V., et al. (2021). Large-scale integration of the plasma proteome with genetics and disease. Nat Genet 53, 1712–1721. 10.1038/s41588-021-00978-w.

41. Burgess, S., Davey Smith, G., Davies, N.M., Dudbridge, F., Gill, D., Glymour, M.M., Hartwig, F.P., Kutalik, Z., Holmes, M.V., Minelli, C., et al. (2019). Guidelines for performing Mendelian randomization investigations: update for summer 2023. Wellcome Open Res 4, 186. 10.12688/wellcomeopenres.15555.3.

42. Hemani, G., Zheng, J., Elsworth, B., Wade, K.H., Haberland, V., Baird, D., Laurin, C., Burgess, S., Bowden, J., Langdon, R., et al. (2018). The MR-Base platform supports systematic causal inference across the human phenome. Elife 7. 10.7554/eLife.34408.

43. Becht, E., McInnes, L., Healy, J., Dutertre, C.A., Kwok, I.W.H., Ng, L.G., Ginhoux, F., and Newell, E.W. (2018). Dimensionality reduction for visualizing single-cell data using UMAP. Nat Biotechnol. 10.1038/nbt.4314.

44. Lange, M., Bergen, V., Klein, M., Setty, M., Reuter, B., Bakhti, M., Lickert, H., Ansari, M., Schniering, J., Schiller, H.B., et al. (2022). CellRank for directed single-cell fate mapping. Nat Methods 19, 159–170. 10.1038/s41592-021-01346-6.

45. Saelens, W., Cannoodt, R., Todorov, H., and Saeys, Y. (2019). A comparison of single-cell trajectory inference methods. Nat Biotechnol 37, 547–554. 10.1038/s41587-019-0071-9.

46. Trapnell, C. (2015). Defining cell types and states with single-cell genomics. Genome Res 25, 1491–1498. 10.1101/gr.190595.115.

47. Yan, D., He, Y., Dai, J., Yang, L., Wang, X., and Ruan, Q. (2017). Vascular endothelial growth factor modified macrophages transdifferentiate into endothelial-like cells and decrease foam cell formation. Biosci Rep 37. 10.1042/BSR20170002.

48. Sayed, N., Wong, W.T., Ospino, F., Meng, S., Lee, J., Jha, A., Dexheimer, P., Aronow, B.J., and Cooke, J.P. (2015). Transdifferentiation of human fibroblasts to endothelial cells: role of innate immunity. Circulation 131, 300–309. 10.1161/CIRCULATIONAHA.113.007394.

49. Takehara, K., LeRoy, E.C., and Grotendorst, G.R. (1987). TGF-beta inhibition of endothelial cell proliferation: alteration of EGF binding and EGF-induced growth-regulatory (competence) gene expression. Cell 49, 415–422. 10.1016/0092-8674(87)90294-7.

50. Tobias, P.S., and Curtiss, L.K. (2007). Toll-like receptors in atherosclerosis. Biochem Soc Trans 35, 1453–1455. 10.1042/BST0351453.

51. Khachigian, L.M. (2023). The MEK-ERK-Egr-1 axis and its regulation in cardiovascular disease. Vascul Pharmacol 153, 107232. 10.1016/j.vph.2023.107232.

52. Pi, S., Mao, L., Chen, J., Shi, H., Liu, Y., Guo, X., Li, Y., Zhou, L., He, H., Yu, C., et al. (2021). The P2RY12 receptor promotes VSMC-derived foam cell formation by inhibiting autophagy in advanced atherosclerosis. Autophagy 17, 980–1000. 10.1080/15548627.2020.1741202.

53. Yoshida, T., Yamashita, M., Horimai, C., and Hayashi, M. (2013). Smooth muscle-selective inhibition of nuclear factor-kappaB attenuates smooth muscle phenotypic switching and neointima formation following vascular injury. J Am Heart Assoc 2, e000230. 10.1161/JAHA.113.000230.

54. Jin, Z., Li, J., Pi, J., Chu, Q., Wei, W., Du, Z., Qing, L., Zhao, X., and Wu, W. (2020). Geniposide alleviates atherosclerosis by regulating macrophage polarization via the FOS/MAPK signaling pathway. Biomed Pharmacother 125, 110015. 10.1016/j.biopha.2020.110015.

55. Akoumianakis, I., Polkinghorne, M., and Antoniades, C. (2022). Non-canonical WNT signalling in cardiovascular disease: mechanisms and therapeutic implications. Nat Rev Cardiol 19, 783–797. 10.1038/s41569-022-00718-5.

56. Fu, X., Sun, Z., Long, Q., Tan, W., Ding, H., Liu, X., Wu, L., Wang, Y., and Zhang, W. (2022). Glycosides from Buyang Huanwu Decoction inhibit atherosclerotic inflammation via JAK/STAT signaling pathway. Phytomedicine 105, 154385. 10.1016/j.phymed.2022.154385.

57. Srinivas, S., Watanabe, T., Lin, C.S., William, C.M., Tanabe, Y., Jessell, T.M., and Costantini, F. (2001). Cre reporter strains produced by targeted insertion of EYFP and ECFP into the ROSA26 locus. BMC Dev Biol 1, 4. 10.1186/1471-213x-1-4.

58. Hassan, H.H., Denis, M., Krimbou, L., Marcil, M., and Genest, J. (2006). Cellular cholesterol homeostasis in vascular endothelial cells. Can J Cardiol 22 *Suppl B*, 35B- 40B. 10.1016/s0828-282x(06)70985-0.

59. Voyta, J.C., Via, D.P., Butterfield, C.E., and Zetter, B.R. (1984). Identification and isolation of endothelial cells based on their increased uptake of acetylated-low density lipoprotein. J Cell Biol 99, 2034–2040. 10.1083/jcb.99.6.2034.

60. Deaton, R.A., Gan, Q., and Owens, G.K. (2009). Sp1-dependent activation of KLF4 is required for PDGF-BB-induced phenotypic modulation of smooth muscle. Am J Physiol Heart Circ Physiol 296, H1027–1037. 10.1152/ajpheart.01230.2008.

61. Cho, D.I., Ahn, M.J., Cho, H.H., Cho, M., Jun, J.H., Kang, B.G., Lim, S.Y., Yoo, S.J., Kim, M.R., Kim, H.S., et al. (2023). ANGPTL4 stabilizes atherosclerotic plaques and modulates the phenotypic transition of vascular smooth muscle cells through KLF4 downregulation. Exp Mol Med 55, 426–442. 10.1038/s12276-023-00937-x.

62. Murgai, M., Ju, W., Eason, M., Kline, J., Beury, D.W., Kaczanowska, S., Miettinen, M.M., Kruhlak, M., Lei, H., Shern, J.F., et al. (2017). KLF4-dependent perivascular cell plasticity mediates pre-metastatic niche formation and metastasis. Nat Med 23, 1176–1190. 10.1038/nm.4400.

63. Chen, Z.Y., Wang, X., Zhou, Y., Offner, G., and Tseng, C.C. (2005). Destabilization of Kruppel-like factor 4 protein in response to serum stimulation involves the ubiquitin-proteasome pathway. Cancer Res 65, 10394–10400. 10.1158/0008-5472.CAN-05-2059.

64. Gamper, A.M., Qiao, X., Kim, J., Zhang, L., DeSimone, M.C., Rathmell, W.K., and Wan, Y. (2012). Regulation of KLF4 turnover reveals an unexpected tissue-specific role of pVHL in tumorigenesis. Mol Cell 45, 233–243. 10.1016/j.molcel.2011.11.031.

65. Libby, P., and Sasiela, W. (2006). Plaque stabilization: Can we turn theory into evidence? Am J Cardiol 98, 26P–33P. 10.1016/j.amjcard.2006.09.017.

66. Brophy, M.L., Dong, Y., Tao, H., Yancey, P.G., Song, K., Zhang, K., Wen, A., Wu, H., Lee, Y., Malovichko, M.V., et al. (2019). Myeloid-Specific Deletion of Epsins 1 and 2 Reduces Atherosclerosis by Preventing LRP-1 Downregulation. Circ Res 124, e6–e19. 10.1161/CIRCRESAHA.118.313028.

67. Dhaliwal, N.K., Miri, K., Davidson, S., Tamim El Jarkass, H., and Mitchell, J.A. (2018). KLF4 Nuclear Export Requires ERK Activation and Initiates Exit from Naive Pluripotency. Stem Cell Reports 10, 1308–1323. 10.1016/j.stemcr.2018.02.007.

68. 68. Yap, C., Mieremet, A., de Vries, C.J.M., Micha, D., and de Waard, V. (2021). Six Shades of Vascular Smooth Muscle Cells Illuminated by KLF4 (Kruppel-Like Factor 4). Arterioscler Thromb Vasc Biol 41, 2693–2707. 10.1161/ATVBAHA.121.316600.

69. Salmon, M., Johnston, W.F., Woo, A., Pope, N.H., Su, G., Upchurch, G.R., Jr., Owens, G.K., and Ailawadi, G. (2013). KLF4 regulates abdominal aortic aneurysm morphology and deletion attenuates aneurysm formation. Circulation 128, S163–174. 10.1161/CIRCULATIONAHA.112.000238.

70. Long, X., Bell, R.D., Gerthoffer, W.T., Zlokovic, B.V., and Miano, J.M. (2008). Myocardin is sufficient for a smooth muscle-like contractile phenotype. Arterioscler Thromb Vasc Biol 28, 1505–1510. 10.1161/ATVBAHA.108.166066.

71. Chattopadhyay, A., Kwartler, C.S., Kaw, K., Li, Y., Kaw, A., Chen, J., LeMaire, S.A., Shen, Y.H., and Milewicz, D.M. (2021). Cholesterol-Induced Phenotypic Modulation of Smooth Muscle Cells to Macrophage/Fibroblast-like Cells Is Driven by an Unfolded Protein Response. Arterioscler Thromb Vasc Biol 41, 302–316. 10.1161/ATVBAHA.120.315164.

72. Zhang, X.H., Zheng, B., Gu, C., Fu, J.R., and Wen, J.K. (2012). TGF-beta1 downregulates AT1 receptor expression via PKC-delta-mediated Sp1 dissociation from KLF4 and Smad-mediated PPAR-gamma association with KLF4. Arterioscler Thromb Vasc Biol 32, 1015–1023. 10.1161/ATVBAHA.111.244962.

73. Tsai, S.Y., Clavel, C., Kim, S., Ang, Y.S., Grisanti, L., Lee, D.F., Kelley, K., and Rendl, M. (2010). Oct4 and klf4 reprogram dermal papilla cells into induced pluripotent stem cells. Stem Cells 28, 221–228. 10.1002/stem.281.

74. Kim, J.B., Zaehres, H., Wu, G., Gentile, L., Ko, K., Sebastiano, V., Arauzo-Bravo, M.J., Ruau, D., Han, D.W., Zenke, M., and Scholer, H.R. (2008). Pluripotent stem cells induced from adult neural stem cells by reprogramming with two factors. Nature 454, 646–650. 10.1038/nature07061.

75. Lee, S., Wottrich, S., and Bonavida, B. (2017). Crosstalks between Raf-kinase inhibitor protein and cancer stem cell transcription factors (Oct4, KLF4, Sox2, Nanog). Tumour Biol 39, 1010428317692253. 10.1177/1010428317692253.

76. Scholer, H.R., Ruppert, S., Suzuki, N., Chowdhury, K., and Gruss, P. (1990). New type of POU domain in germ line-specific protein Oct-4. Nature 344, 435–439. 10.1038/344435a0.

