## Supplemental Figures and Tables 1 and 2 for "Epsins oversee smooth muscle cell reprograming by influencing master regulators KLF4 and OCT4"

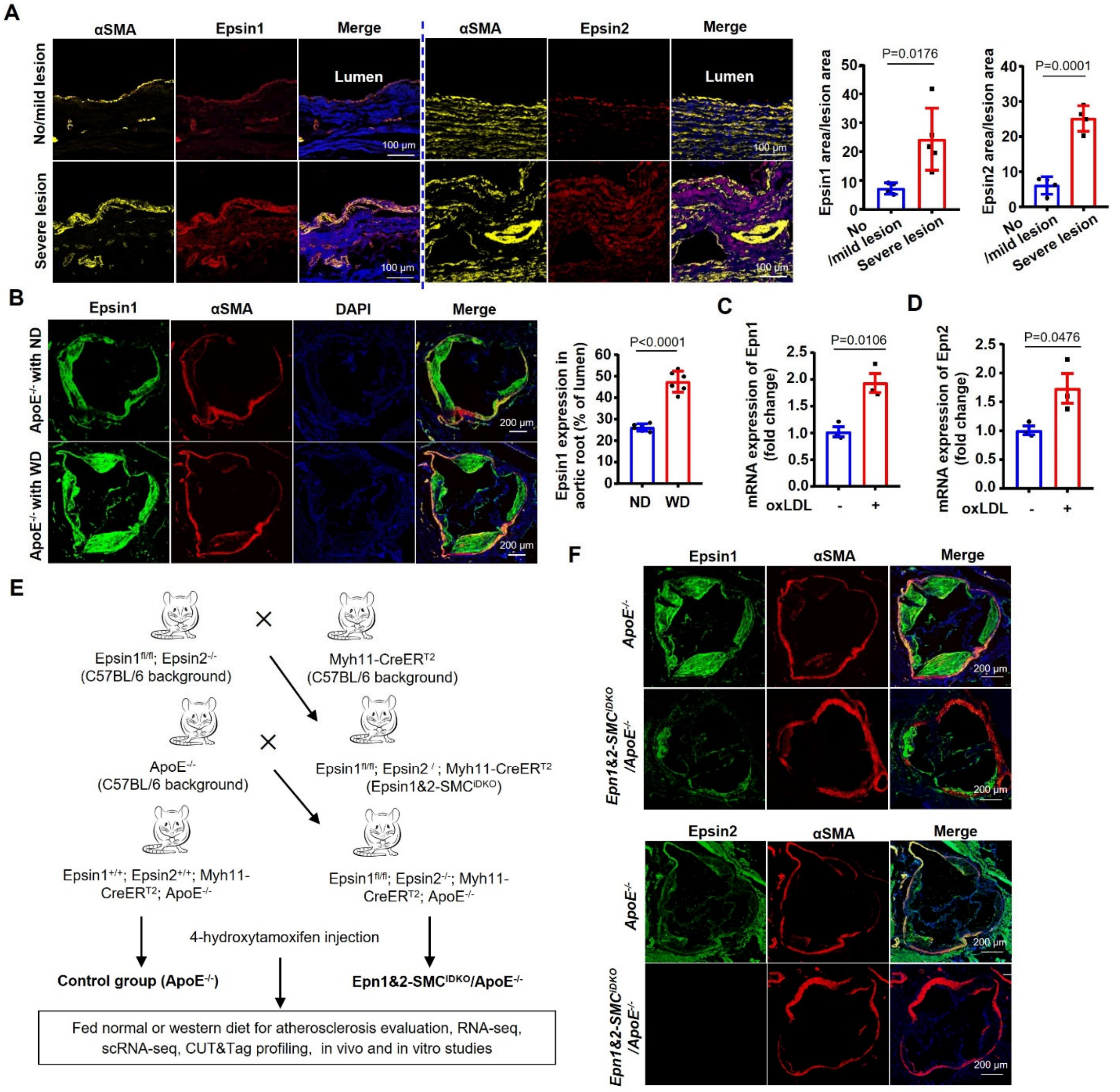


**Figure S1. Upregulated Expression of Epsins in VSMCs in Response to Atherosclerotic Stimuli**, **related to Figures 1 and STAR Methods.**
(A) Immunofluorescence staining for Epsin1, Epsin2, α-SMA in aortae from human patients with no/mild or severe atherosclerotic lesions. Scale bar=100 µm. n=4-5 samples. (B) Immunofluorescence staining of Epsin 1 and α-SMA in *ApoE*$^{-/-}$ mice fed ND or WD. Scale bar=200 µm. Quantitation of Epsin1 expression in aortic roots. n=6 mice. (C-D) Transcript abundance of Epsins1 (C) and Epsin2 (D) in SMCs after 12 hrs exposure to 100 µg/mL oxLDL. n=3 independent repeats. e, Strategy for generation of mouse models. (F) Immunofluorescence staining for Epsin 1, Epsin 2 and α-SMA in aortic root of *ApoE*$^{-/-}$ and *Epn1&2-SMC*$^{iDKO}$*/ApoE*$^{-/-}$ mice fed a WD for 16 weeks. SMC, aortic smooth muscle cell; EC, endothelial cell; WD, western diet; ND, normal diet. All *P* values were calculated using two-tailed unpaired Student's *t*-test. Data are mean ± s.d.

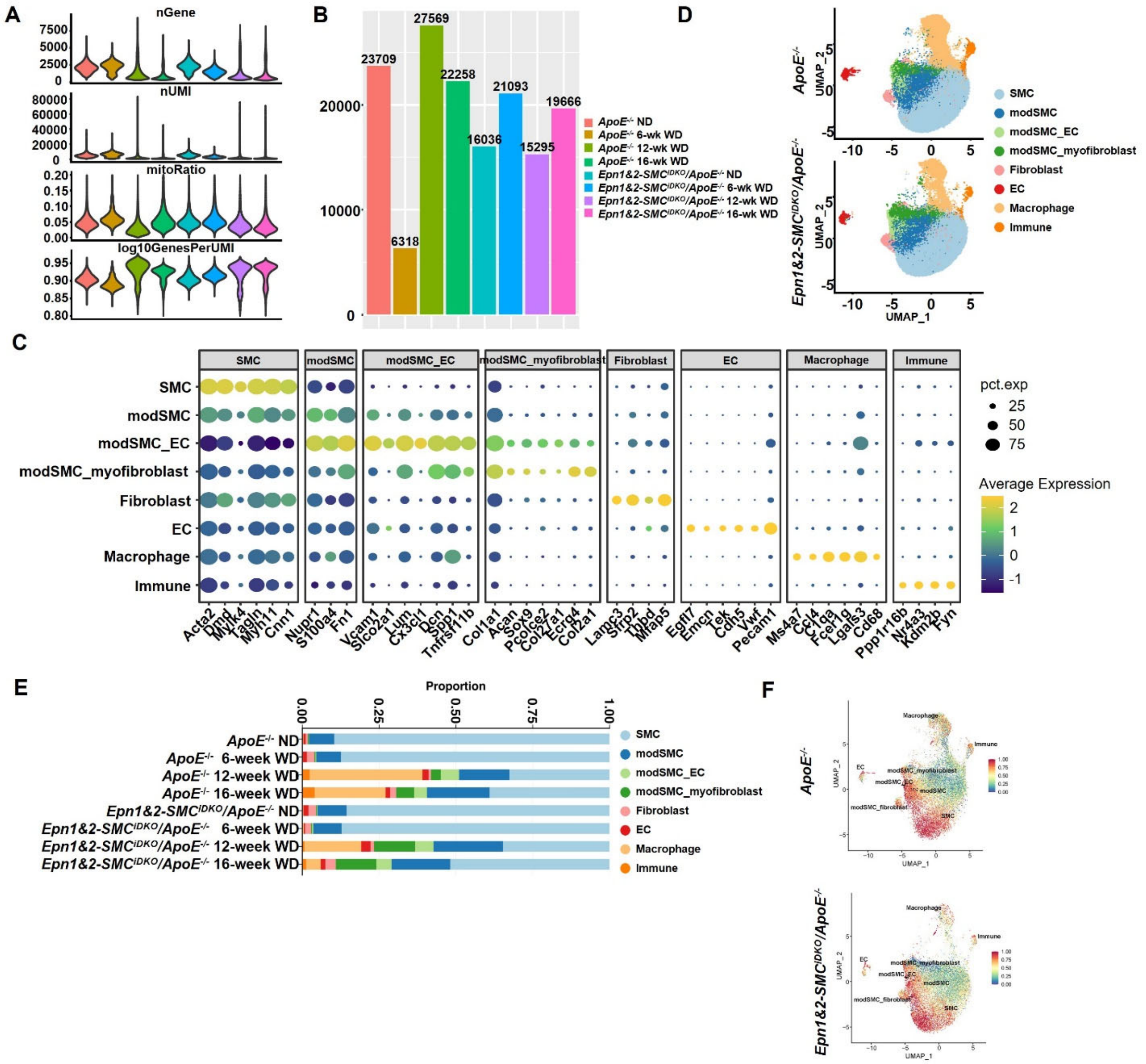

**Figure S2. Single-cell Transcriptomic Profiling of Aortae from Atherosclerotic Mice, related to Figures 1.**

(A-B) Quality control of scRNA-seq data in mouse model. Violin plot (A) of the number of genes and UMI, the mitochondrial ratio, and gene/UMI across each mouse model. Bar plot (B) of available cells from various mouse models across different diet feeding-week. (C) Dot heatmap showing the expression of typical marker genes for major cell clusters in mouse models. (D) UMAP visualization of major cell clusters across each mouse model. (E) Stacked bar plot showing the proportion of major cell clusters derived from mouse models across various diet feeding-week. (F) UMAP visualization of inferred CytoTRACE scores for major cell types between *ApoE*$^{-/-}$ and *Epn1&2-SMC*$^{iDKO}$*/ApoE*$^{-/-}$ mice. UMAP, Uniform Manifold Approximation and Projection.

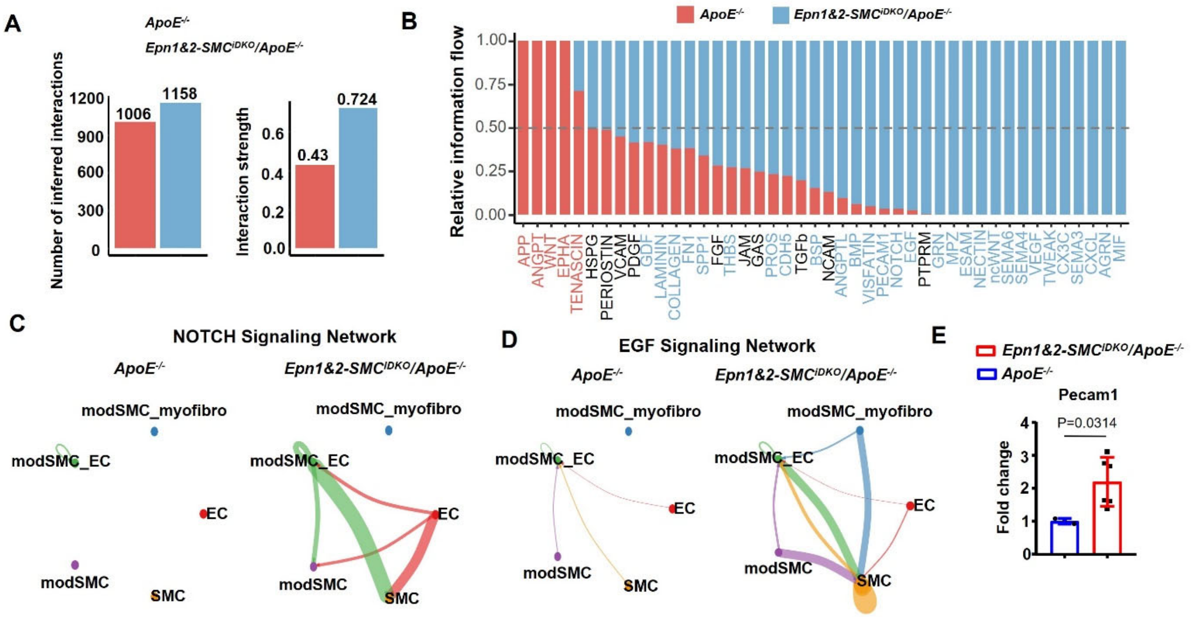


**Figure S3. Cell-to-cell Communications analysis of scRNA-seq Data, related to Figures 1.**
(A) Global summary of number and strength of cell-to-cell interactions among cell clusters in aortae of *ApoE*$^{-/-}$ and *Epn1&2-SMC*$^{iDKO}$*/ApoE*$^{-/-}$ mice, identified using *CellChat*. (B) SMC transition-relevant signaling pathways modeluated by SMC-specific Epsins deficiency, identified using *CellChat*. (C-D) Cell communication networks of NOTCH (C) and EGF (D) signaling across SMC phenotypic modulation between *ApoE*$^{-/-}$ and *Epn1&2-SMC*$^{iDKO}$*/ApoE*$^{-/-}$ mice. (E) The mRNA levels of EC marker *Pecam1* was determined in aortae of *ApoE*$^{-/-}$ and *Epn1&2-SMC*$^{iDKO}$*/ApoE*$^{-/-}$ mice fed a WD for 16 weeks. *P* values were calculated using two-tailed unpaired Student's *t*-test. Data are mean ± s.d. n=3-6 mice. SMC, aortic smooth muscle cell; EC, endothelial cell; WD, western diet.

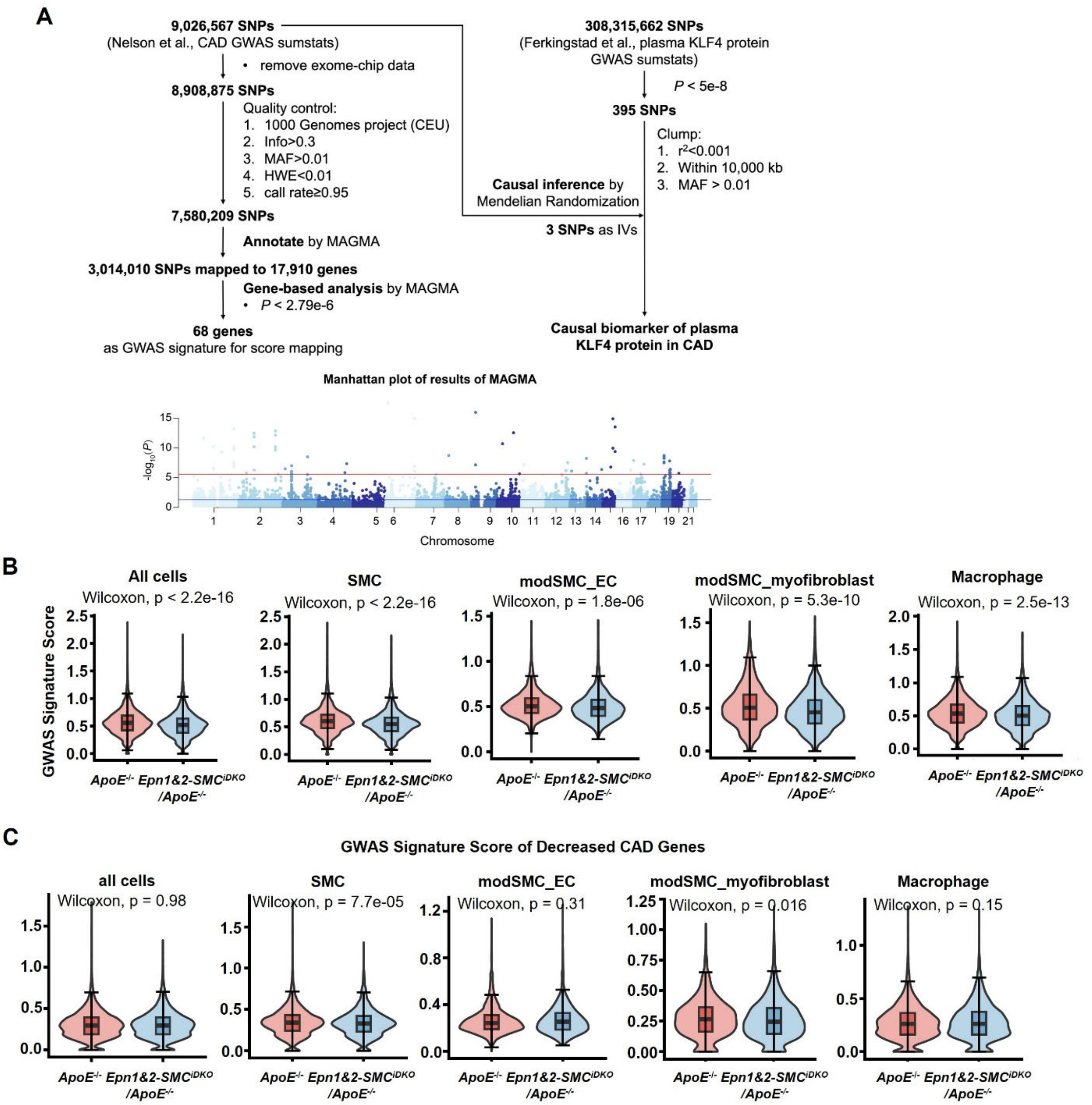


**Figure S4. Mapping of CAD-associated genes identified by GWAS in mouse aortae scRNA-seq Data, related to Figures 2.**

(A) Flowchart of the identification of CAD susceptibility genes and the causal inference of plasma KLF4 on CAD risk. Manhattan plot showing the genome-wide susceptibility genes of CAD using MAGMA. The red line represents the genome-wide significance at $P$<0.05/17910. (B-C) Differential signature score of GWAS-identified CAD-associated genes between *ApoE*$^{-/-}$ and *Epn1&2-SMC*$^{iDKO}$/*ApoE*$^{-/-}$ mice across major cell types from scRNA-seq data. $P$ value was calculated using Wilcoxon rank sum test. 19 of the 68 CAD susceptibility genes are associated with increased CAD risk (B) and 30 with decreased CAD risk (C). SMC, aortic smooth muscle cell; GWAS, genome-wide associated studie; CAD, coronary artery disease. Data are mean ± s.d.

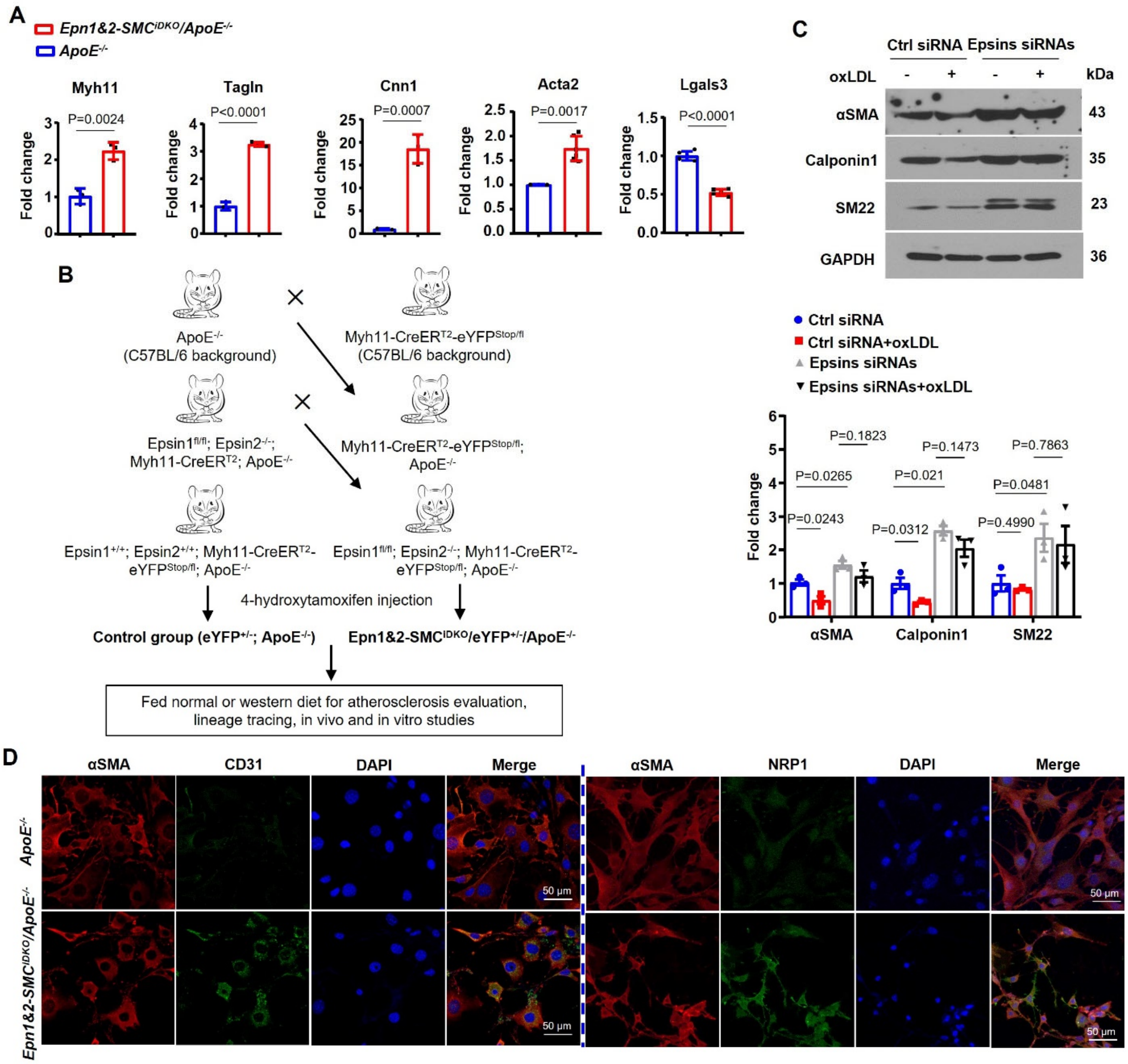


**Figure S5. SMC Differentiation Markers are Regulated by Epsins during Atherosclerosis, related to Figures 2, 3 and STAR Methods**.

(A) Transcript abundance of SMC marker genes (*Myh11*, *Tagln*, *Cnn1* and *Acta2*) and macrophage marker *lgals3* in aortae of *ApoE*$^{-/-}$ and *Epn1&2-SMC*$^{iDKO}$*/ApoE*$^{-/-}$ mice fed a western diet (WD) for 16 weeks. n=3-6 mice. (B) Breeding scheme to establish the compound mutant mouse strains. (C) Immunoblot of primary SMCs transfected with control siRNA or siRNAs against Espins 1&2 followed by stimulation with 100 μg/mL oxLDL with antibodies against SMC markers (α-SMA, Calponin1 and SM22). n=3 independent repeats. (D) Immunofluorescence staining of long-term cultured SMC isolated from *ApoE*$^{-/-}$ and *Epn1&2-SMC*$^{iDKO}$*/ApoE*$^{-/-}$ mice with or without the treatment of 100 μg/mL oxLDL for 48 hrs with antibodies against. EC markers CD31/NRP1 and α-SMA. Scale bar=50 μm. SMC, aortic smooth muscle cell; EC, endothelial cell; WD, western diet; siRNA, small interfering RNA; oxLDL, oxidized low-density lipoprotein. All *P* values were calculated using two-tailed unpaired Student's *t*-test. Data are mean ± s.d.

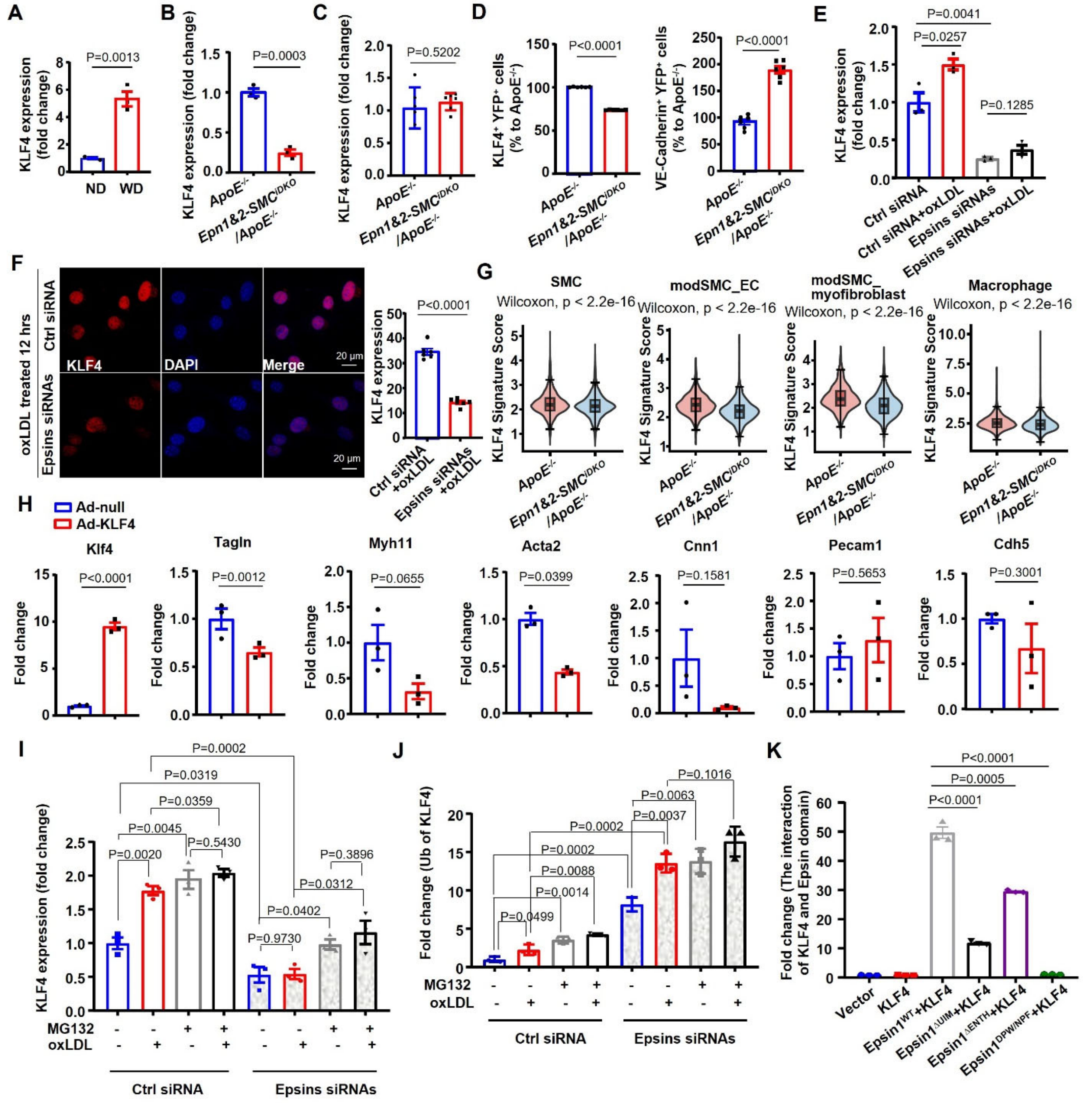

**Figure S6. Epsin Suppresses Expression of SMC markers by Decreasing KLF4 Expression, related to Figures 4.**

(A-B) Quantification of KLF4 protein level in aortae from *ApoE*$^{-/-}$ mice fed a ND or WD for 16 weeks (A) or from *ApoE*$^{-/-}$ and *Epn1&2-SMC*$^{iDKO}$*/ApoE*$^{-/-}$ mice (B) fed a WD for 16 weeks described in Figure 4C,D. n=3 mice. c, Transcript abundance of KLF4 in total cells dissociated from the aortae of *ApoE*$^{-/-}$ and *Epn1&2-SMC*$^{iDKO}$*/ApoE*$^{-/-}$ mice fed a WD for 16 weeks. n=3 mice. (D) Quantification of the number of VE-Cadherin$^{+}$ and KLF4$^{+}$ cells in YFP$^{+}$ cells sorted from the aortae of *YFP*$^{+/-}$*/ApoE*$^{-/-}$ and *Epn1&2-SMC*$^{iDKO}$*/YFP*$^{+/-}$*/ApoE*$^{-/-}$ mice fed on WD for 14 weeks described in Figure 4g. n=6 mice. e, Quantification of the KLF4 protein levels in SMCs transfected with control siRNA or Epsin 1&2 siRNAs following treatment with 100 μg/mL oxLDL described in Figure 4i. n=3 independent repeats. (F) Immunofluorescence staining of KLF4 in primary SMCs isolated from the aortae of *ApoE*$^{-/-}$ and *Epn1&2-SMC*$^{iDKO}$*/ApoE*$^{-/-}$mice after treatment with 100 μg/mL oxLDL for 12 hrs. Scale bar=20 μm. n=6 independent repeats. (G) Differential signature score of KLF4 binding genes in major aortae cell types from *ApoE*$^{-/-}$ and *Epn1&2-SMC*$^{iDKO}$*/ApoE*$^{-/-}$ mice derived from scRNA-seq data. *P* value was calculated by Wilcoxon rank sum test. The signature score was calculated using differential 1113 target genes of KLF4 binding site with *PercentageFeatureSet* function deposited in *Seurat*. (H) The mRNA levels of *KLF4*, SMC marker genes and EC markers were determined in primary SMCs from *Epn1&2-SMC*$^{iDKO}$*/ApoE*$^{-/-}$ mice transfected with Ad-null or Ad-KLF4 for 24 hrs, followed by 12 hrs oxLDL treatment. n=3 independent repeats. (I-J) Quantification of western blot analyses of KLF4 expression (I) and ubiquitination (J) in SMCs transfected with control siRNA or Epsin 1&2 siRNAs following treatment with 100 nM MG132 or 100 μg/mL oxLDL described in Figure 4k. n=3 independent repeats. (K) Quantification of the immunoblot result of the interaction between the domain of Epsin1 and KLF4 of Figure 4l. n=3 independent repeats. SMC, aortic smooth muscle cell; EC, endothelial cell; WD, western diet; siRNA, small interfering RNA; oxLDL, oxidized low-density lipoprotein; Ad, adenovirus. All *P* values were calculated using two-tailed unpaired Student's *t*-test except (G). Data are mean ± s.d.

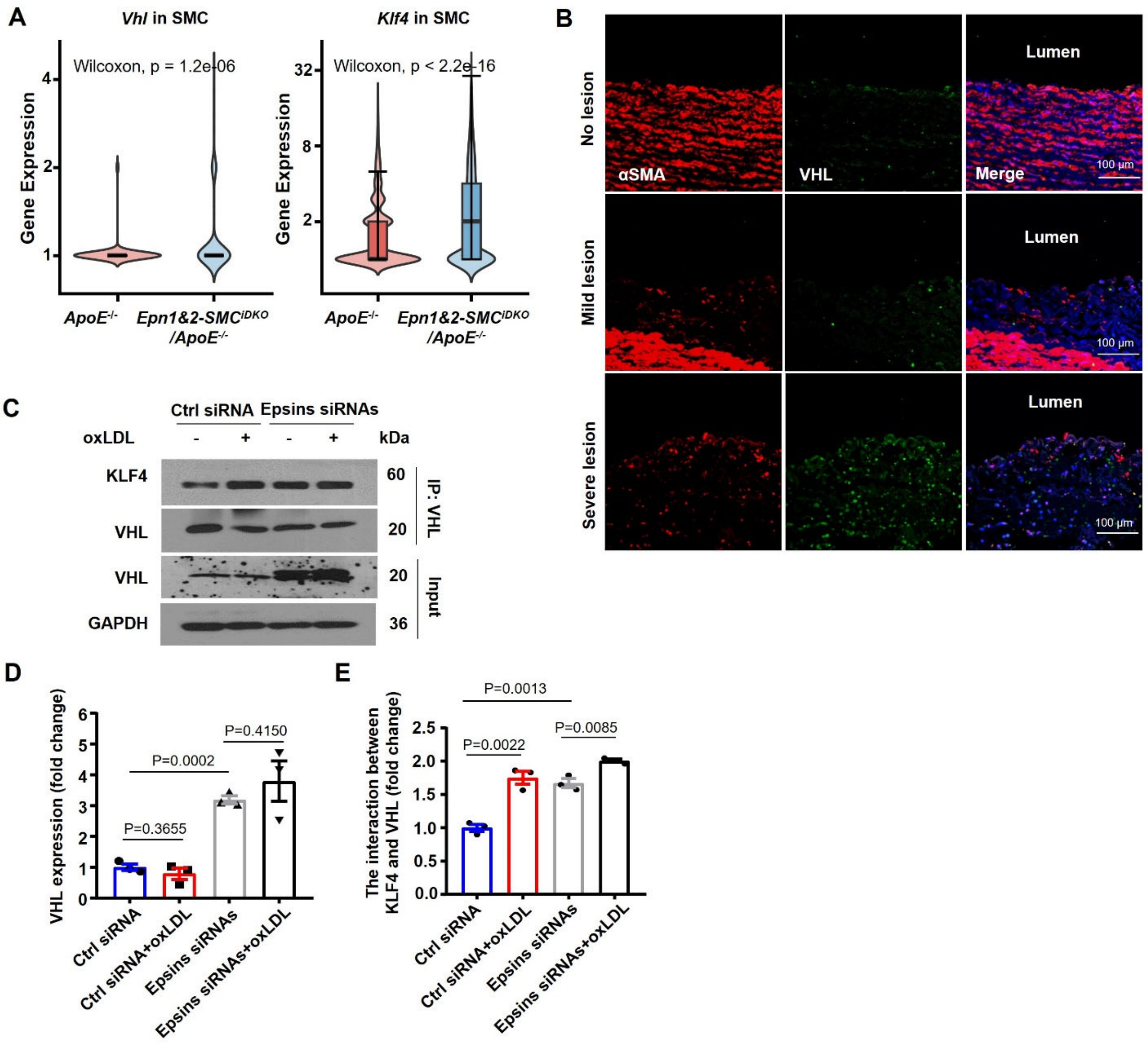


**Figure S7. Epsins Hinder KLF4 Ubiquitination by Preventing VHL Binding to KLF4, related to Figures 4.**

(A) Differential gene expression of *Vhl* and *Klf4* in SMC cells from aortae of *ApoE*$^{-/-}$ and *Epn1&2-SMC*$^{iDKO}$*/ApoE*$^{-/-}$ mice derived from scRNA-seq data. *P* value was calculated by Wilcoxon rank sum test. (B) Immunofluorescence staining for α-SMA, VHL of aortae from human patients with no, mild, or severe atherosclerotic lesions. Scale bar=100 μm. (C-E) Immunoprecipitation of VHL and KLF4 with anti-VHL antibody in primary SMCs isolated from *ApoE*$^{-/-}$ and *Epn1&2-SMC*$^{iDKO}$*/ApoE*$^{-/-}$mice with or without oxLDL (100 μg/mL) treatment and analyzed with western blot. *P* values were calculated using two-tailed unpaired Student's *t*-test except g. Data are mean ± s.d. n=3 independent repeats. SMC, aortic smooth muscle cell; EC, endothelial cell; oxLDL, oxidized low-density lipoprotein.

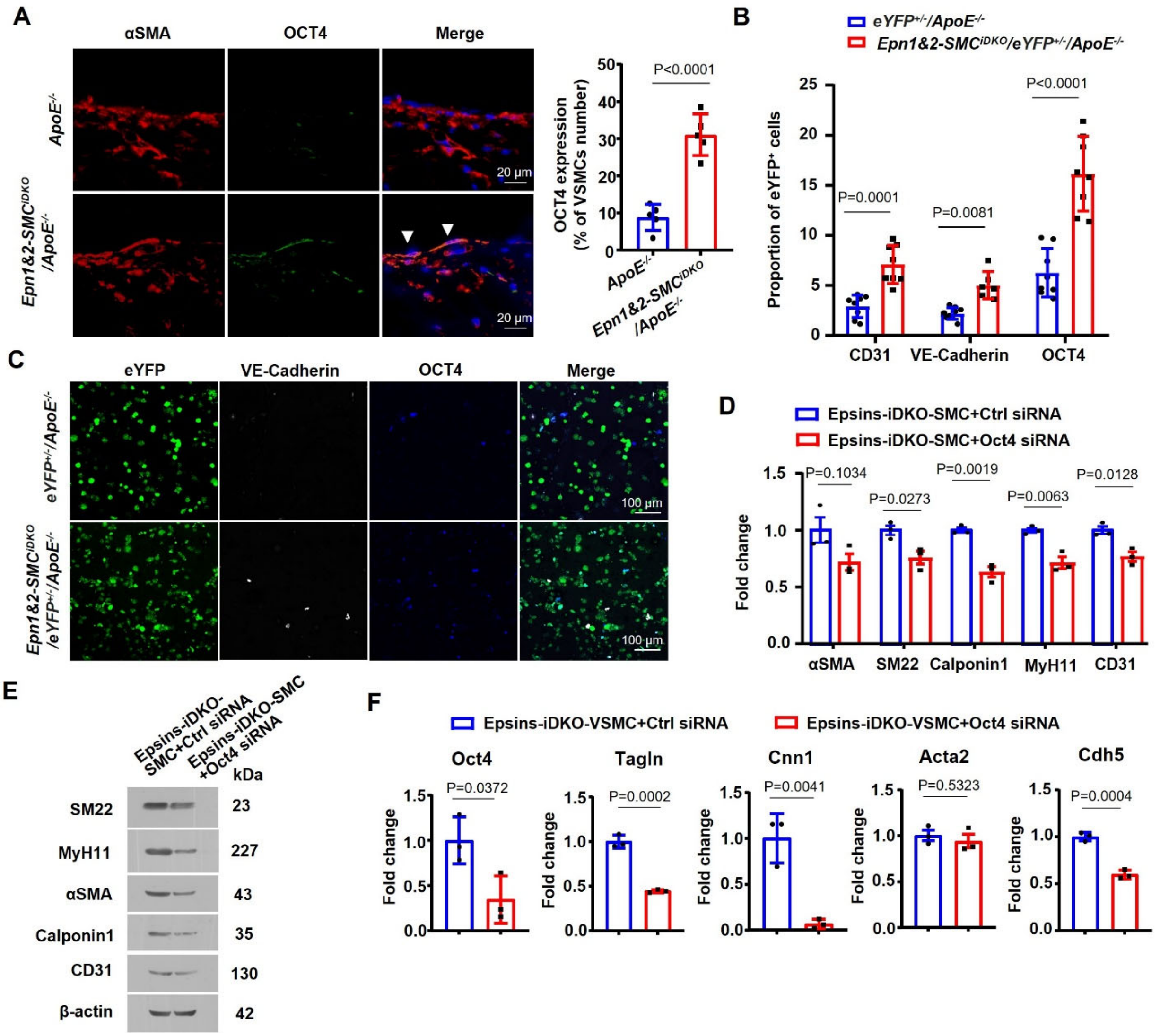


**Figure S8. SMC-specific Epsin Deficiencies Augments OCT4 Expression in Atherosclerotic Plaques, related to Figures 5.**

(A) Immunofluorescence staining for α-SMA and OCT4 in aortic roots of *ApoE*$^{-/-}$ and *Epn1&2-SMC*$^{iDKO}$*/ApoE*$^{-/-}$ mice fed a WD for 16 weeks. Scale bar=20 µm. n=5 mice. (B-C) The localization of OCT4 and VE-Cadherin expression in YFP-tagged cells sorted from *YFP*$^{+/-}$*/ApoE*$^{-/-}$ and *Epn1&2-SMC*$^{iDKO}$*/YFP*$^{+/-}$*/ApoE*$^{-/-}$ mice fed a WD for 16 weeks, as revealed by confocal microscopy analysis. Scale bar=100 µm. (B) Quantitation of the proportion of CD31$^{+}$, VE-Cadherin$^{+}$ and OCT4$^{+}$ in YFP$^{+}$ cells. n=7 mice. (D-F) The protein (D-E) and mRNA (F) levels of SMC differentiation markers and EC markers were measured in long-term cultured SMCs from *Epn1&2-SMC*$^{iDKO}$*/ApoE*$^{-/-}$ mice transfected with control or Oct4 siRNA for 48 hrs and quantitation. n=3 independent repeats. *P* values were calculated using two-tailed unpaired Student's *t*-test. Data are mean ± s.d. SMC, aortic smooth muscle cell; EC, endothelial cell; siRNA, small interfering RNA; WD, western diet.

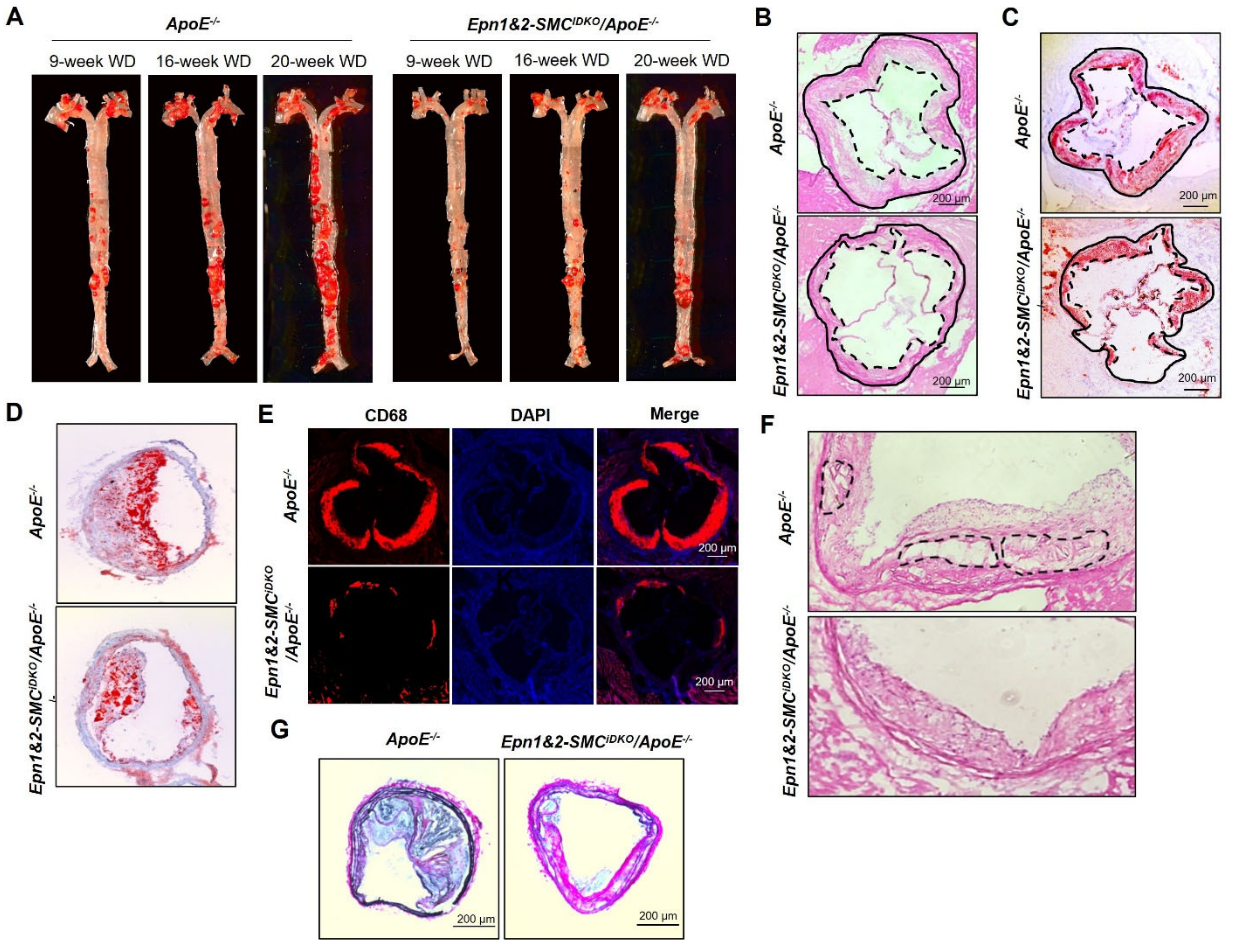


**Figure S9. SMC-specific Epsin 1 and 2 Deficiency Reduces Atherosclerotic Plaques and Enhances Stability of Lesions in Mice, related to Figures 6.**

*ApoE*$^{-/-}$ and *Epn1&2-SMC*$^{iDKO}$*/ApoE*$^{-/-}$ mice were fed a WD for 9, 16 and 20 weeks. The sections of aortic root and brachiocephalic trunks were collected from the *ApoE*$^{-/-}$ and *Epn1&2-SMC*$^{iDKO}$*/ApoE*$^{-/-}$ mice fed a WD for 16 weeks. (A) *En face* Oil Red O staining of the whole aortas. (B) Hematoxylin and eosin staining of aortic roots showed the size of atherosclerotic lesion. (C-D) Oil Red O staining of aortic roots (C) and brachiocephalic trunks (D) were used to show the lipid accumulation in the lesion. (E) Immunofluorescence staining of CD68 in aortic roots were performed to show the inflammation in lesion. (F) Hematoxylin and eosin staining of aortic roots showed the necrotic cores in the lesion. Necrotic cores were outlined in black dash. (G) Verhoeff-Van Gieson's staining of brachiocephalic trunks was performed to show the stability of the lesion. Scale bar=200 mm. SMC, aortic smooth muscle cell; EC, endothelial cell; WD, western diet.

**Table S1. Antibodies used in this study**

| Antibody | Sources | Cat# | Source/ Isotype | Applications | Cross activity | MW (kDa) |
|---|---|---|---|---|---|---|
| KLF4 | Abcam | ab129473 | Rabbit | WB, ICC/IF | H M | 60 |
| OCT4 | Abcam | ab200834 | Rabbit | WB, IHC-P, ICC/IF, IP, F | H M | 45 |
| VHL | Cell signaling | 68547S | Rabbit | WB | H | 18~22 |
| Ubiquitin | Cytoskeleton, Inc | AUB01 | Mouse | WB, IF | M | |
| Cleaved-caspase3 | Cell signaling | 9661 | Rabbit | WB, IHC, IF, IP, F | H M R Mk | 17 19 |
| NRP1 | Abcam | ab81321 | Rabbit | WB, IHC-P, ICC/IF, IP, F | H M R | 103 |
| Galectin3 | Abcam | ab2785 | MS | WB, ICC/IF, IHC-P | H M | 26 |
| Ecrg4 | Abcam | ab224077 | MS | WB, IHC-P | H | 17 |
| α-SMA-cy3 | Sigma | C6198-.2ml | MS | IF | H M | |
| VE-Cadherin | R&D | AF1002 | Goat | WB, IHC | M | |
| MYH11 | Abcam | ab224804 | Rabbit | IF, WB | M R H | 42 |
| SM22 (Tagln) | Abcam | ab14106 | Rabbit | ICC/IF, WB | M R H | 23 |
| Calponin 1 | Abcam | ab46794 | Rabbit | WB, IHC-P, ICC/IF | M R H P | 34 |
| CD31 | BD Pharmingen | 550274 | Rat | IF | M H R | |
| CD68 | Santa Cruz | sc-20060 | MS | IF | M H | |
| Epsin 1 | Home made | n/a | Rabbit | WB, IP, IF | H M | 75~80 |
| Epsin 2 | Home made | n/a | Rabbit | WB, IP, IF | H M | 65~70 |
| HA.11 | BioLegend | 901502 | MS | WB, IP | all | n/a |
| ICAM-1 | Santa Cruz | sc-8419 | MS | WB, IP, IF, IHC(P), F | M H R | |
| P-selectin | Santa Cruz | sc-271267 | MS | WB, IP, IF, ELISA, F | M H R | |
| ß-actin | Cell signaling | 4967 | Rabbit | WB | H M R, etc | 45 |
| GAPDH | Santa Cruz | Sc-166545 | MS | WB | H M | 37-43 |

| Secondary antibodies (Invitrogen) | | |
|---|---|---|
| **Alexa Fluor 488 (Green)** | **Alexa Fluor 594 (Red)** | **Alexa Fluor 647 (Blue)** |
| Donkey anti-Mouse (A21202) | Donkey anti-Mouse (A23744) | Donkey anti-Mouse (A31571) |

| | | |
|---|---|---|
| Donkey anti-Rabbit (A21206) | Donkey anti-Rabbit (A21207) | Donkey anti-Rabbit (A31573) |
| Donkey anti-Rat (A21208) | Donkey anti-Rat (A21209) | Goat anti-Rat (A21247) |
| Donkey anti-Goat (A11055) | Donkey anti-Goat (A11058) | |

**Table S2. Primers used in the RT-PCR for mouse EndoMT marker genes**

| Mouse genes | Primers |
|---|---|
| **KLF4** | KLF4_L1 5'-GATCTCGGGCAATCTGGGG-3'<br>KLF4_R1 5'- TTCCTCACGCCAACGGTTAGT-3' |
| **Cdh5 (VE-Cadherin)** | VE-Cadherin_L1 5'-TACCACTTCAAGCTGCCAGA-3'<br>VE-Cadherin_R1 5'-TCGGAAGAATTGGCCTCTGT-3' |
| **PECAM-1 (CD31)** | PECAM-1_L1 5'-GGTCGTGAATGACACCCAAG-3'<br>PECAM-1_R1 5'-ACTCTGACTGCAAGAGTGCT-3' |
| **Oct4** | Oct4_L1 5'-AAGTTGGCGTGGAGACTTT-3'<br>Oct4_R1 5'-TTCCACCTTCTCCAACTTCAC-3' |
| **Acta2 (α-SMA)** | Acta2_L1 5'-GTCCCTCTATGCCTCTGGAC-3'<br>Acta2_R1 5'-AAGGAATAGCCACGCTCAGT-3' |
| **Myh11** | Myh11_L1 5'- CTCTGGCCTCTTCTGTGTGG-3'<br>Myh11_R1 5'-TTTCCAGCTCCCCCGTGAT-3' |
| **Cnn1** | Cnn1_L1 5'-CTGTTGCGCTTGTCTGTGTC-3'<br>Cnn1_R1 5'-TGCAGCTTGTTGATAAATTCGC-3' |
| **Tagln** | Tagln_L1 5'-GTGAGCCAAGCAGACTTCCAT-3'<br>Tagln_R1 5'-CCAATTTGCTCAGAATCACACC-3' |
| **Lgals3** | Lgals3_L1 5'-CTAATCAGGAAAATGGCAGACAG-3'<br>Lgals3_R1 5'-TCCTTGAGGGTTTGGGTTTC-3' |
| **Ecrg4** | Ecrg4_L1 5'-AGATGCTCCAGAAACGAGAAG-3'<br>Ecrg4_R1 5'-TCCTTTGCTGTGTTCTCGG-3' |
